# How the eyes move, not where they land, predicts what we remember across the lifespan

**DOI:** 10.64898/2026.08.14.744828

**Authors:** Iryna Schommartz, Bhavin Choksi, Gemma Roig, Benjamin de Haas, Yee Lee Shing

## Abstract

Where and how we move our eyes through a natural scene depends jointly on the scene and on the viewer. How the spatial and temporal organization of viewing changes across the lifespan, and whether those changes relate to memory, remains unclear. We recorded eye movements from a lifespan cohort (N = 179, ages 5–79) during free viewing of naturalistic scenes, then tested recognition across graded levels of image degradation. Characterizing each observer by how closely their viewing corresponded to that of age peers, young adults, and a stimulus-driven salience model, we found a developmental dissociation: consistency in where the eyes were directed increased monotonically with age, whereas consistency in how they moved — saccade direction, length, and fixation duration — followed an inverted-U peaking in young adulthood. Recognition sensitivity followed an inverted-U of the same form. Across all three reference frames, typicality in how the eyes moved, but not in their spatial targeting, predicted recognition.

## Introduction

When we encounter a scene — a zoo entrance, a street corner, a cluttered kitchen — our gaze follows an idiosyncratic path through it. Whether a five-year-old, a young adult, or their grandparent, each observer fixates a somewhat different slice of the same visual world, reflecting both the pull of scene content and stable individual differences in oculomotor behavior^1–6^. This variation is not inconsequential: gaze exploration during encoding is causally linked to memory formation. Restricting eye movements at encoding impairs subsequent recognition memory^7,8^, and the number and duration of fixations made during free viewing scales directly with hippocampal engagement and subsequent memory^9–12^. What supports memory, however, appears to lie not only at where we look but also in the sequential how — the temporal architecture of visual sampling^13–15^. Gaze reinstatement studies show that the spatial overlap between encoding and retrieval fixations predicts recognition^16,17^, and that it is specifically the sequential fidelity with which eye movements from encoding are re-enacted at retrieval — how fixations unfold in order and time, rather than merely which locations are revisited — that predicts the quality of episodic recollection^18,19^. This work, however, concerns the correspondence between encoding and retrieval gaze, and treats the encoding scanpath as a single entity. Whether the spatial and dynamic components of encoding gaze itself contribute differentially to what is later remembered — independent of retrieval reenactment — remains unknown.

Importantly, the findings above were based almost entirely on young adult samples. However, we know that human oculomotor behavior follows an ontogenetic trajectory: scanning behavior matures through childhood and adolescence^1,20–22^, plateaus in young adulthood, and shifts with senescence in ways that may reflect either strategic adaptation or declining neural integrity^23–25^, or neurological insult^26–28^. The gaze-memory relationship itself appears to change with age: older adults show altered fixation distributions, execute more similar eye movements across repeated presentations, and the coupling between spatial gaze sampling and subsequent recognition is disrupted^29,30^. At the other end of the lifespan, children’s fixation patterns are more idiosyncratic and less canonical, becoming adult-like only through adolescence^20^. Whether this developmental immaturity carries consequences for memory encoding — analogous to the disruptions observed in aging — has never been directly tested. Nor is it known whether maturation and senescence affect all properties of viewing alike: gaze has often been indexed globally, as a single canonicality score or as spatial overlap, leaving open whether the trajectories of where and how the eyes move diverge. Memory itself, moreover, is not equally robust across the lifespan at the point of retrieval: everyday recognition rarely operates on a repetition of the complete experienced past, and recovering a memory from a partial cue requires pattern completion — a hippocampally-dependent process that matures through childhood and is vulnerable in ageing^31–33^. Testing recognition under graded visual degradation is therefore a good way to probe the resilience of memory across the lifespan, not merely whether encoding succeeded.

Answering this requires a reference against which individual and developmental differences can be measured. Classical saliency models predict gaze from low-level features^34,35^, or their integration with semantic content^36,37^, and deep convolutional networks have substantially improved fixation prediction by incorporating scene context^38–40^. These models remain fundamentally spatial, however: they predict where fixations land, not the order in which they unfold. A recent class of scanpath models addresses this by generating fixation sequences rather than static salience maps. DeepGaze III^41^ is a current state of the art: it conditions each successive fixation on the observer’s recent oculomotor history, so that scanpath history constrains where sampling goes next over and above scene content alone. Trained on young-adult free-viewing data, it serves as a descriptive reference for how closely an individual’s scanpath matches a stimulus-driven sampling pattern — not as a normative standard of correct viewing.

Here we aimed to test where age-related divergence in scene viewing originates, and whether it carries consequences for memory. We recorded eye movements from 179 participants ages 5 to 79 (grouped into five age groups for analyses) during free viewing of 60 naturalistic scenes, followed later by a cued-recognition task in which studied scenes and semantic lures appeared at six graded levels of image degradation, separating baseline recognition sensitivity from resilience to visual challenge. We quantified viewing with MultiMatch^42^, which compares any two eye-movement trajectories along four independent properties: the spatial locations fixated, and the direction, amplitude, and timing with which gaze moved between them. MultiMatch is comparative: it quantifies not the observer’s viewing in isolation, but how closely it corresponds to a reference trajectory, such that a high value on a given dimension means the observer moved their eyes as the reference did along that property. We therefore characterized each observer by their typicality relative to three references: same age peers, indexing consistency within a cohort; young adults, indexing conformity to a mature pattern; and DeepGaze III, a stimulus-driven saliency model trained on adult free-viewing data, indexing agreement with image-driven sampling. We fitted quadratic age trajectories to each dimension, characterized recognition sensitivity across degradation levels, and tested whether typicality along any dimension predicted subsequent recognition, controlling for image memorability.

We found a developmental dissociation. Peer consistency in where the eyes were directed increased monotonically with age, whereas consistency in how they moved — saccade direction and length and duration — followed an inverted-U, peaking in young adulthood. Typicality in how the eyes moved, but not where they landed, predicted recognition across all three references.

## Results

### Natural scene gaze consistency separates into a stimulus-driven and an individual component

To quantify how much of scene-viewing consistency is attributable to the stimulus and how much to the observer, we compared the scanpaths of every pair of observers viewing the same image, across the full sample of 179 participants (aged 5-79; **Fig.1d**). We aggregated similarity to the focal observer (mean similarity to all other observers on that image; **Fig. 1a-c**) and partitioned in into variance attributable to the focal observer (Agent) — the stable tendency of an observer to be similar to others across scenes, independent of what they viewed — variance attributable to the image (Image), and residual observer-by-image variation (**Fig. 2a**). The same decomposition was applied to DeepGaze III scanpaths^41^. Because the model samples scanpaths probabilistically from a scene-driven distribution rather than from individual observers, two model agents viewing the same image differ only by sampling noise, not by identity; this image-driven stochastic variation is the level of agent variance expected in the absence of any individual signature and therefore provides the floor against which human agent variance is compared.

**Figure 1.**
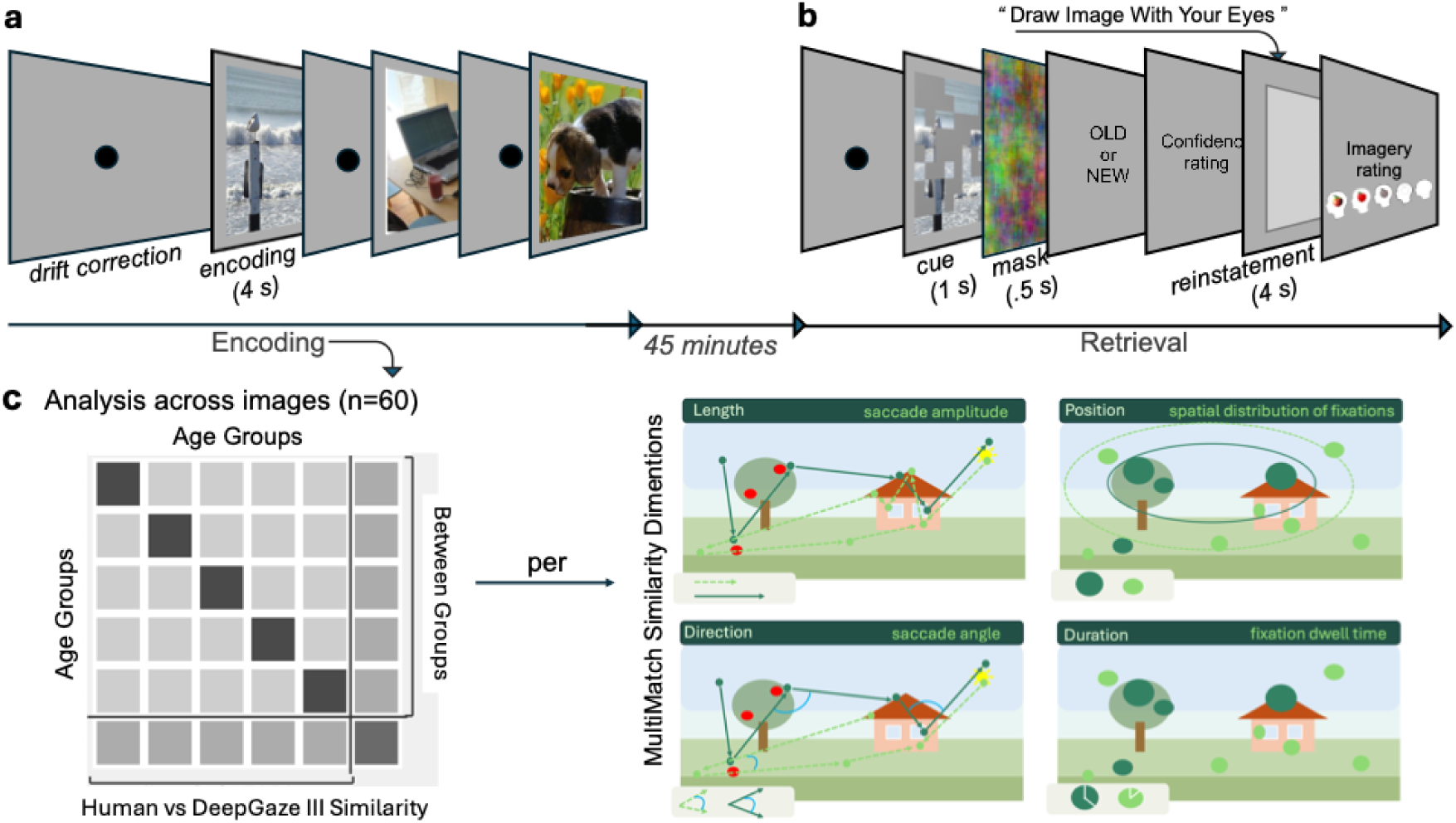
Task design, MultiMatch similarity dimensions, and analysis structure. **a**, Encoding phase. Each trial began with a central drift-correction target that remained until fixated (response-terminated), re-calibrating the gaze-position estimated before stimulus onset. Participants then freely viewed 60 naturalistic scenes for 4 s each while eye movements were recorded. Stimuli were drawn from the OSIEshort set ^43^ of the OSIE dataset ^44^. **b**, Retrieval phase, administered approximately 45 minutes after encoding. Each trial began with a 1 s probe cue under varying degrees of occlusion (30–80%), followed by a 0.5 s mask, after which participants judged whether the cue image was old or new, reported recognition confidence using a five-point scale, and were instructed to reinstate the image by tracing it with their eye movements (“Draw image with your eyes”) for 4 seconds, followed by a second imagery confidence rating. Probe images comprised the original 60 studied scenes and 60 semantic lures. **c**, MultiMatch similarity dimensions and analysis structure. Four independent dimensions quantify distinct spatiotemporal properties of the scanpath: Length (saccade amplitude; solid = canonical scanpath, dashed = divergent), Position (spatial distribution of fixations; circle size = dwell time), Direction (saccade angle) and Duration (fixation dwell time distribution). The group similarity matrix (averaged across 60 images) has rows and columns for the five age groups (5–7, 8–10, 11–12, 19–30, 65–80 years) and the DeepGaze III agent (DG III); diagonal cells show within-group similarity; off-diagonal cells between-group similarity; and the bottom row/rightmost column human–DG III similarity.

**Figure 2.**
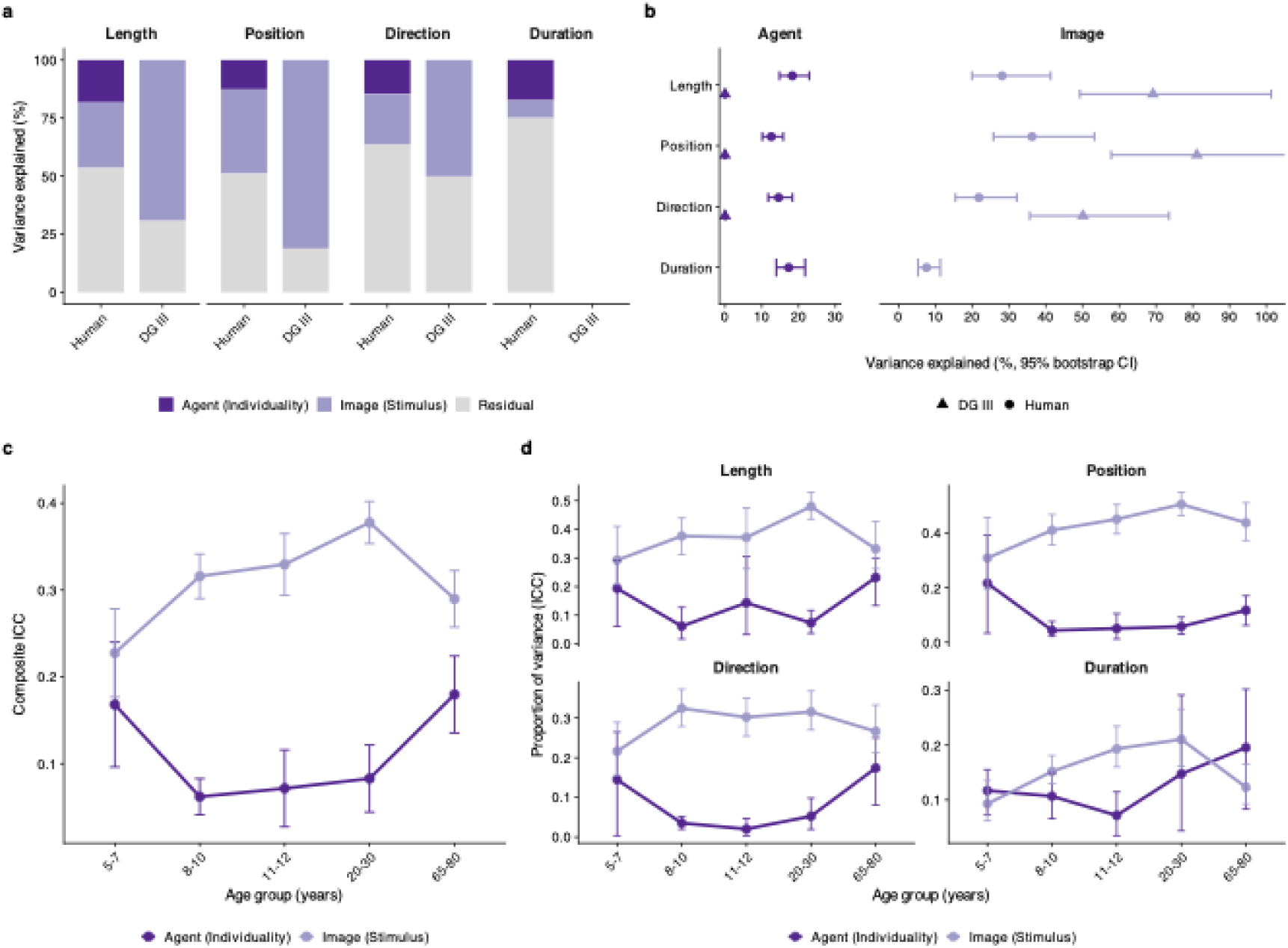
Variance decomposition of scanpath similarity and its developmental trajectory across the lifespan. **a,** Scanpath similarity along four dimensions, partitioned by linear mixed-effects models with crossed random intercepts for observer and image and no fixed effects. Stacked bars show variance attributable to the observer (deep purple; idiosyncratic signature), the image (light purple; stimulus-dependency), and the residual (grey), which principally reflects observer-by-image variation. Duration is undefined for DeepGaze III, which assigns every fixation an identical duration. **b,** Agent and Image components with 95% bootstrap confidence intervals; note the differing x-axis ranges. Circles, humans; triangles, DeepGaze III. **c,** Composite trajectory, averaging each component across the four dimensions within each age group. Error bars are approximate 95% confidence intervals, pooling the per-metric bootstrap standard errors (the dimensions share observers and images); the weighted mixed models reported in the text are the primary analysis. **d,** Within each age group, variance in pairwise similarity among same-age observers was partitioned into a participant-level component (ICC_Agent,_) and an image-level component (ICC_Image_). Markers are means across 1,000 bootstrap resamples of observers; error bars are 95% bootstrap confidence intervals. Free y-axis scales show the relative developmental pattern within each dimension. Residual variance is not shown (Supplementary Fig. S1).

In humans, similarity was shaped jointly by the stimulus and the observer: image properties accounted for 7.6–36.3% of total variance across dimensions, and stable individual signatures for a further 12.6–18.3% (largely independent of age group; Supplementary Table S3-4; **Fig. 2a-b**; the complete ICC breakdown with CIs). In DeepGaze III the balance shifted decisively toward the stimulus: image properties accounted for 49.9–79.8% of variance, while agent-level variance was indistinguishable from zero on every dimension (all ICC < 0.07), as its architecture predicts. Against this baseline, human agent variance reflects idiosyncratic structure over and above stimulus-driven salience. Human image-driven variance was 28.0–43.5 percentage points lower than the model’s on every dimension, the remainder carried by the observer and by observer-specific responses to scenes (**Fig. 2b**). Human gaze is therefore far less tightly governed by stimulus salience than a purely image-driven process.

Fixation duration could not be compared, as DeepGaze III assigns every fixation an identical duration and so produces no temporal variation to decompose. In humans, duration carried the strongest individual signature of any dimension (17.3%) alongside the weakest stimulus contribution (7.6%), making it the most observer-driven and least image-driven property we measured — whereas the model was image-dominated throughout.

### Gaze composition shifts non-linearly across the lifespan

To examine how individual gaze signatures change across the lifespan, we estimated the Agent and Image components separately within each age group and dimension, bootstrapping over observers (1,000 iterations), and modelled the cohort-level estimates as a function of age (Methods). The allocation of variance shifted non-linearly with age (individual differences reliable in every cell, all *P* < 0.001, Supplementary Table S5; **Fig. 2c**). Idiosyncrasy (ICC_Agent_) followed a U-shaped trajectory (quadratic *b* = 0.18, *t* = 5.94, *P* < 0.001), highest in childhood and older adulthood and lowest in young adulthood, while stimulus-dependency (ICC_Image_) showed the mirror pattern (quadratic *b* = −0.17, *t* = −5.06, *P* < 0.001) peaking in young adulthood. Because the components sum to one by construction, these describe a single reallocation between individual signature and stimulus control rather than two independent effects. This reallocation was uneven across dimensions (**Fig. 2d**): Agent and Image followed reliably different age trajectories for Direction (component × age interaction, *F*_(2,4)_ = 13.3, *P* = 0.017) and Duration (*F*_(2,4)_ = 10.1, *P* = 0.027), and marginally for Length and Position (*P* = 0.062 and 0.064). Component-wise, idiosyncrasy curved reliably only for Direction (quadratic *P* = 0.014) and stimulus-dependency only for Duration (*P* = 0.045). The direction of the effect was consistent across all four dimensions, with agent variance lowest in young adults and elevated at both extremes.

### Kinematic and temporal gaze consistency change most across the lifespan

Having established that the composition of gaze variance shifts across the lifespan, we next asked how consistent viewing is within each age group, and along which dimensions that consistency changes across the lifespan. For each MultiMatch dimension we modelled within-group scanpath similarity — the similarity of each observer to their same-age peers — as a function of age group.

Within-group similarity differed reliably across age groups on every dimension, but to very different degrees, with the largest differences for Duration similarity (*F*_(4,174)_ = 20.86, *P* < 0.001, *ω^2^_p_* = 0.31), followed by Direction (*F*_(4,174)_ = 11.00, *P* < 0.001, *ω^2^_p_* = 0.18), Length (*F*_(4,174)_ = 8.42, *P* < 0.001, *ω^2^_p_* = 0.14), and Position (*F*_(4,173)_ = 3.63, *P* = 0.007, *ω^2^_p_* = 0.06; **Fig. 3a**). We decomposed each dimension’s ordered group means into orthogonal linear and quadratic components (Methods). Direction and Duration similarity were each dominated by a negative quadratic component, with no linear trend (Direction: quadratic *t*_(174)_ = –6.48, *p* = 9.0×10⁻¹□, *ω^2^_p_* = 0.19, linear *P* = 0.56; Duration: quadratic *t*_(174)_ = –7.75, *P* = 7.4×10⁻^13^, *ω^2^_p_* = 0.23, linear p = 0.19) — a symmetric inverted-U peaking in young adulthood, confirmed by reliable rise and fall contrasts (all *P* < 0.001; Supplementary Table S6). Length similarity showed both a positive linear (*t*_(174)_ = 4.58, *P* = 9.0×10⁻□, *ω^2^_p_* = 0.10) and a weaker negative quadratic component (*t*(_174)_ = −2.88, *P* = 0.004; *ω^2^_p_* = 0.04): consistency rose into young adulthood and then largely plateaued (fall contrast *P* = 0.055), the linear increase dominating. Position similarity showed the opposite profile — a positive linear trend (*t*_(174)_ = 3.16, *P* = 0.002, *ω^2^_p_*= 0.04) and no quadratic component (*P* = 0.999) — a monotonic increase in spatial consistency across the lifespan, with the youngest children the least consistent and older adults the most (the youngest children differed reliably only from older adults, 5-7 vs 65-80 *P* = 0.002; Supplementary Tables S6-S7).

**Figure 3.**
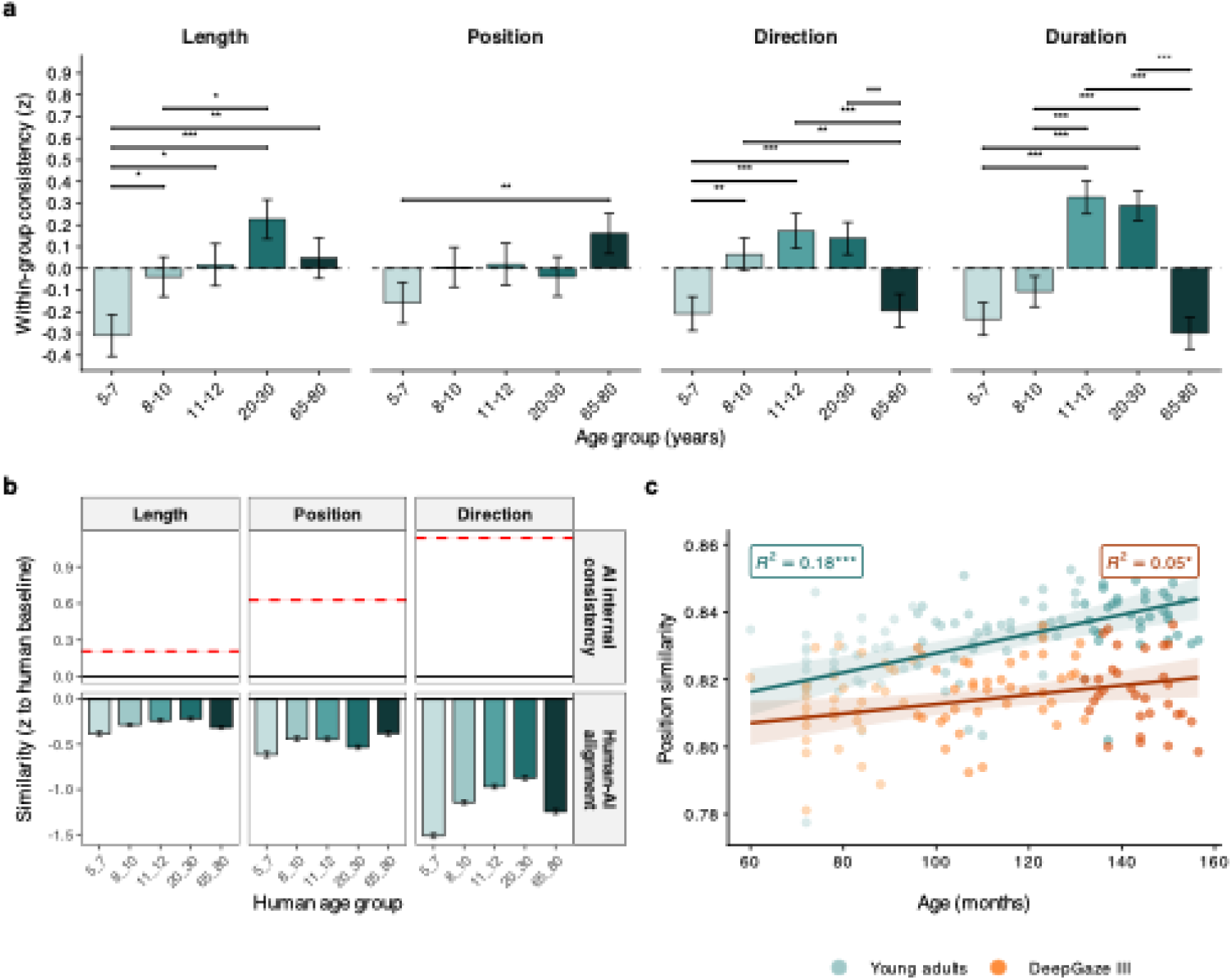
Within-group scanpath consistency and its relation to a stimulus-driven model across the lifespan. **a,** Within-group similarity for four MultiMatch dimensions across five age groups. Each dimension is z-scored on its own distribution (y = 0 is its mean within-group similarity and bars show relative departure from that average). Bars are estimated marginal means from linear mixed-effects models with crossed random intercepts for observer and image; error bars are ±1 model-based s.e.m. Brackets show all significant Tukey-adjusted pairwise contrasts (*P < 0.05; **P < 0.01; ***P < 0.001); the full matrix is in Supplementary Table S7. **b,** Alignment of human scanpaths to DeepGaze III for Length, Position and Direction (duration is undefined for DeepGaze III), z-scored to the human– human baseline (z = 0, solid black line); this baseline differs from that in **a** (see Methods), so values are not directly comparable across panels. Top row: model-internal consistency, the deterministic ceiling of DeepGaze III self-similarity (red dashed line, mean ± s.e.m.). Bottom row: human–DeepGaze III alignment by age group (bars, mean; error bars ±1 s.e.m.). **c,** Children’s spatial targeting matures toward adults, not the model. Each child’s Position similarity to individual young adults (teal) and DeepGaze III (orange) against age in months; points colored by child age group, lines are linear fits with 95% CI. Other dimensions in Supplementary Fig. S3.

Together these results reveal a dissociation in the shape of developmental change. Consistency in the temporal and kinematic dimensions changed non-monotonically, rising to a young-adult peak and declining in older age — sharply for fixation duration and saccade direction, and more weakly for saccade length. Spatial-position consistency, by contrast, increased early in development, and stayed plateaued across the lifespan. Where the eyes are sent thus becomes steadily more consistent with age, whereas how they move follows an inverted-U — the dynamic dimensions are the locus of the strongest, and the only non-monotonic, developmental change.

### Humans share scanpath structure with each other but not with the model

To test whether human gaze carries systematic structure beyond image-driven salience, we compared scanpath similarity across three pairing types — human–human, human–model (DeepGaze III), and model–model — with crossed random intercepts for agent and image (**Fig. 3b**). Pairing type modulated similarity on every dimension (Length *ω^2^_p_* = 0.09; Position *ω^2^_p_* = 0.26; Direction *ω^2^_p_* = 0.56; all *P* < 0.001). Model–model pairs were the most similar, but this reflects the model’s construction: because its agents are independent probabilistic samples from a single scene-driven distribution, they differ only in how dispersed that distribution is for each image, not by any individual identity. The informative comparison is therefore between the two cross-source pairings, and here human observers resembled one another more than they resembled the model on every dimension (all *P* < 0.001). Because DeepGaze III’s spatial predictions target group-level fixation tendencies rather than individual scanpaths, a comparison against individual humans is conservative for the model on spatial position; a reference against a representative-group (per-image consensus) young adult yields the same conclusion (below; Supplementary Table S7-S8). Even so, human observers resembled one another more than the model on every dimension, most strongly on the dynamic dimensions the model is not built to capture.

### Children develop their gaze toward human, not model, structure

If development carries gaze toward specifically human structure, children should approach the adult pattern more and faster than that of the model. Regressing children’s similarity to young adults, and to DeepGaze III, on age in months (controlling for sex, n = 103), similarity to adults increased with age on every dimension (Length *b* = 7.9×10⁻□, Position *b* = 2.9×10⁻□, Direction *b* = 3.0×10⁻□, Duration *b* = 4.1×10⁻□; all *P* ≤ 0.001), and similarity to the model increased more slowly. For Position, children converged on young adults roughly twice as fast as on the model (adults *b* = 2.9×10⁻□, *t* = 4.89, *P* < 0.001; model *b* = 1.4×10⁻□, *t* = 2.38, *P* = 0.019), a reliable difference in rate (reference × age interaction *b* = −1.5×10⁻⁴, *t* = −5.33, *P* < 0.001; **Fig. 3c**). The spatial effect held against a per-image consensus young adult — the medoid young-adult scanpath, a representative-group reference — (consensus − model *b* = 2.7×10⁻□, *t* = 6.42, *P* < 0.001; Supplementary Table S7-8). Development thus carries children’s gaze toward spatial regularities of adult scanning that a salience-trained model does not reproduce. In older adults, neither reference varied with age on any dimension (all *P* > 0.48; Supplementary Table S10).

### Seeding the model with human fixations does not recover the missing structure

The preceding analyses leave open why the model misses human structure. One possibility is that DeepGaze III views each scene without human context, so seeding it with an observer’s own initial fixations might close the gap. We tested this by seeding the model with each participant’s first three, five or ten fixations and comparing the resulting human–model similarity to the free-viewing baseline (**Fig. 4**). Seeding did not bring the model’s scanpaths closer to human gaze on any dimension. Positional similarity was unchanged across seeding levels (*F*_(3,1039)_ = 2.03, *P* = 0.11, , *ω^2^_p_* = 0.003), and although directional and length similarity varied statistically (Direction *F*_(3, 1051)_ = 11.8, *P* < 0.001, *ω^2^_p_* = 0.03; Length *F*_(3, 1031)_ = 22.8, *P* < 0.001, *ω^2^_p_* = 0.06), the changes were negligible in magnitude — under 0.006 similarity units from no seeding to ten seeded fixations, if anything slightly toward human gaze rather than away. These effects did not vary with age on any dimension (all age x seeding interactions *ω^2^_p_* ≍ 0, *P* > 0.99). Providing the model with a human’s own fixation history therefore neither improves nor degrades its match to the human scanpaths in any substantive way — the structure the model fails to reproduce is not information it could be handed at the start of a trial, but structure of a kind it does not represent.

**Figure 4.**
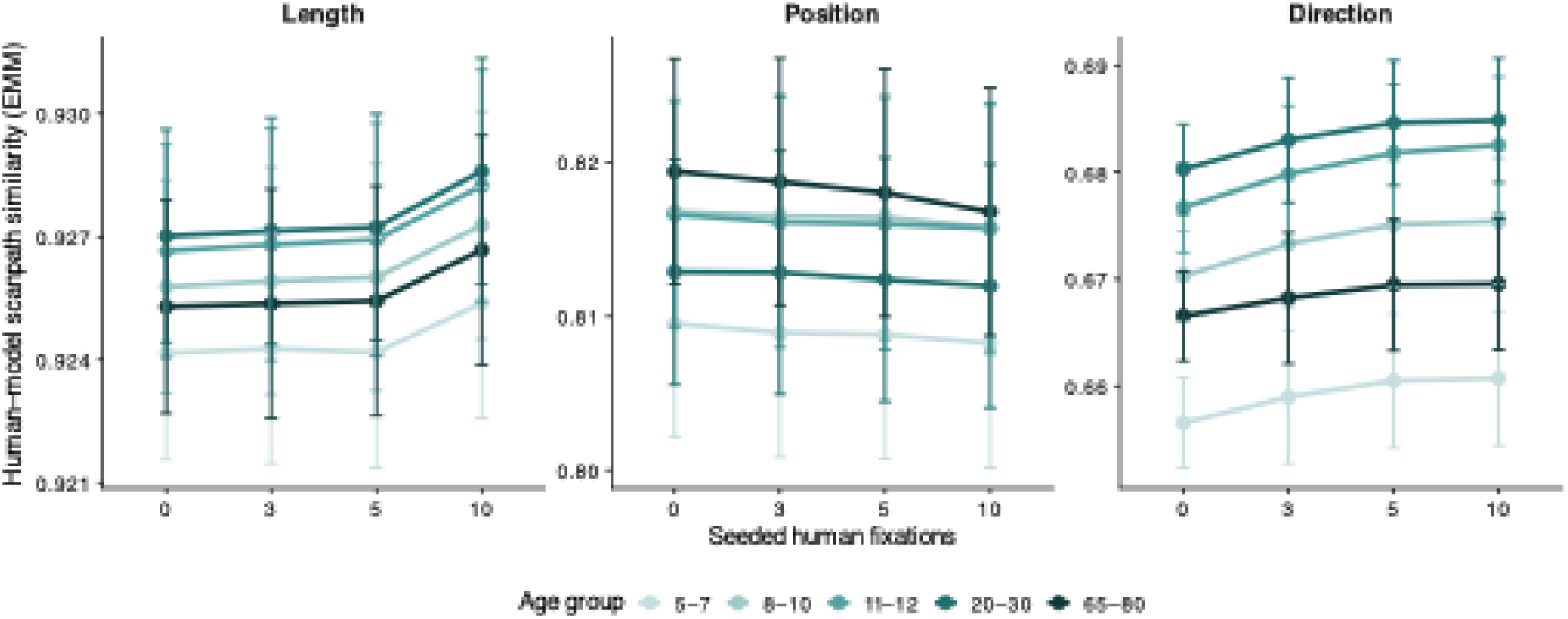
Seeding DeepGaze III with human fixations does not move its scanpaths toward human gaze. Estimated marginal means (± 95% CI) of human–model scanpath similarity as a function of the number of human fixations used to seed the model (0 = free viewing, 3, 5, 10), shown separately for saccadic length, spatial position, and saccadic direction, and by age group (light-to-dark teal across the lifespan). Means are from linear mixed-effects models with age group × seeding level as fixed effects and crossed random intercepts for participant and image, fitted separately per dimension.

### An inverted U-trajectory of recognition, robust to visual degradation

Having characterized how visual exploration matures and where it departs from a stimulus-driven model, we turn to one of its functional consequences: we first establish how recognition develops across the lifespan, then ask whether encoding gaze predicts what is later recognized. Successful recognition of degraded input depends on pattern completion – reconstructing a stored representation from partial cues. To characterize recognition and its robustness to visual challenge across the lifespan, we measured recognition sensitivity (d′) across graded image occlusion (0.3-0.8), fitting a linear mixed model with occlusion as a continuous predictor, a second-order polynomial for age, and image memorability (ResMem; Needell & Bainbridge, 2022) as a covariate, with random intercepts for participants.

Recognition sensitivity followed an inverted-U across the lifespan (quadratic age term *b* = –8.98, *t_(_*_196)_ = –7.03, *P* < 10^−10^, ω *^2^_p_* = .22, continuous age; group-midpoint coding gave the same result, *b* = −8.68, *t*_(198)_ = –6.75, *P* < 10^−9^, ω*^2^_p_* = .21, Supplementary Table S11): performance rose steeply through childhood, plateaued from late childhood through adulthood, and declined in older age (**Fig. 5a**). Image memorability had a small independent effect on recognition (*b* = 0.05, *P* = 0.005, ω*^2^_p_* = 0.007). Occlusion reduced recognition in a threshold-like manner — preserved across mild degradation, falling in discrete steps at higher levels (Supplementary Table S12, **Fig. S4a**) — yet did not reshape the developmental trajectory: neither the quadratic age × occlusion interaction (*P* = 0.59) nor in a fully categorical age-group × occlusion interaction (*P* = 0.22) was significant. The inverted-U held across all levels of visual challenge, age groups remaining at near-constant distances as occlusion increased. Tukey-adjusted contrasts localized the trajectory: the youngest children (5–7) were outperformed by every older group (all *P* < .001); performance then plateaued (8–10, 11–12, and 20–30 mutually indistinguishable, all *P* > .09); older adults, outperformed by each of these (all *P* ≤ .002), performed at the level of the youngest children (vs 5–7, *P* = 0.84), completing the inverted-U (Supplementary **Fig. S4b;** Supplementary Table S13).

**Figure 5.**
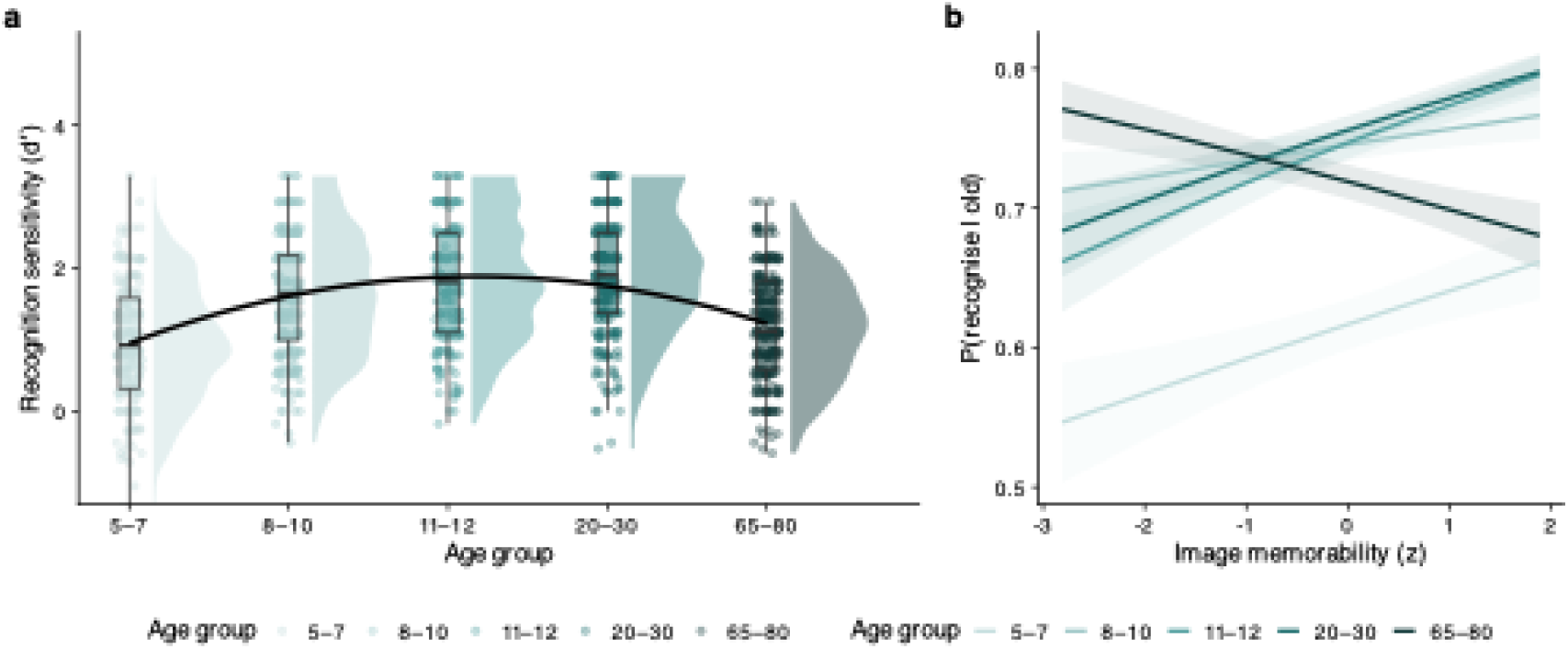
Recognition sensitivity across the lifespan and under visual degradation. **a**, Recognition sensitivity (d′) by age group, pooled across occlusion levels; raincloud plots show the distribution, individual participants (points), and interquartile range (boxplots), with a second-order polynomial fit (black line). Performance rose through childhood, plateaued in late childhood and adulthood, and declined in older age — an inverted-U (quadratic age term P < 0.001, continuous-age model; full pairwise contrasts in Supplementary Table S13). b, Old-item recognition accuracy as a function of intrinsic image memorability (ResMem), by age group; lines show model-predicted slopes with 95% CI. Age group as in the legend.

Age groups also differed in how they used intrinsic image memorability. In a trial-level model of old-item recognition, the memorability benefit varied with age: younger groups showed a positive relationship between an image’s ResMem memorability and recognition, whereas in older adults this benefit was abolished (memorability × 65–80, interaction *b* = −0.20, *z* = −2.79, *P* = 0.005; older-adult simple slope *b* = −0.11, 95% CI [−0.29, 0.07]; Supplementary Table S14; **Fig. 5b**). Older adults thus failed to capitalize on the stimulus-intrinsic properties that aided younger observers — a memory-domain parallel to their reduced reliance on stimulus-driven salience during viewing. This reflects a genuine loss of the memorability benefit rather than false recognition of memorable-looking lures, whose memorability did not predict older adults’ false alarms (b = 0.01, P = 0.92; Supplementary Table S17).

### Scanpath dynamics, not spatial targets, predict recognition

Finally, we asked whether gaze and memory are linked: does the typicality of an individual’s scanpath predict whether they later recognize what they saw? We tested three references — similarity to age-matched peers, to the young-adult prototype, and to DeepGaze III — modelling trial-level recognition of studied items as a function of encoding-scanpath typicality on all four MultiMatch dimensions, with image memorability, occlusion, and image and participant identity controlled (**Fig.7**).

**Figure 7.**
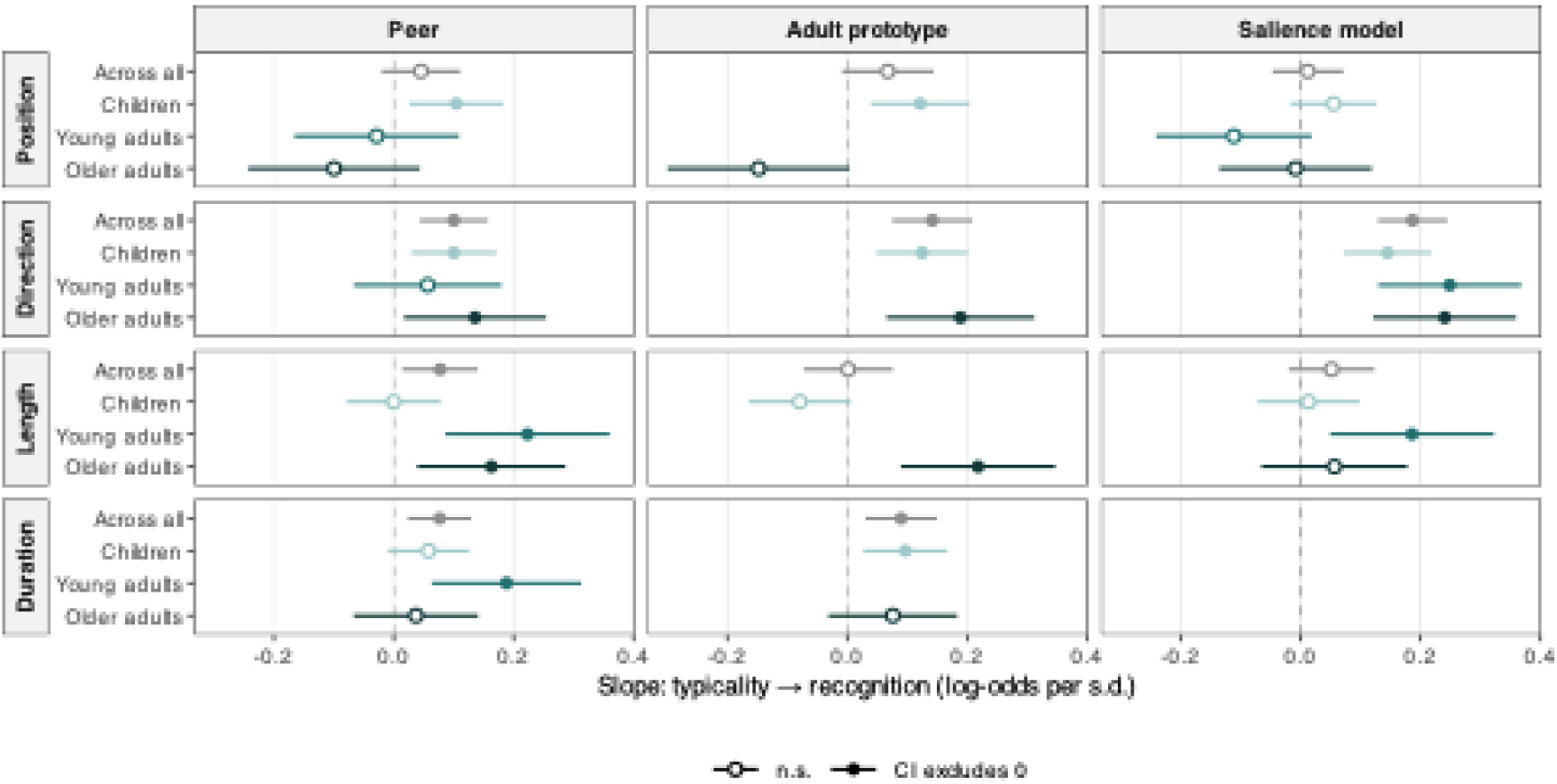
Gaze typicality predicts recognition, dimension-specifically and with age-dependent structure. Slopes relating the typicality of a participant’s encoding scanpath to subsequent recognition of that scene (log-odds of a correct old-item response per +1 s.d. of similarity), from mixed-effects logistic models with image memorability, occlusion, and crossed random intercepts for participant and image. Columns are the three reference templates against which typicality was measured — same-age peers, the young-adult prototype, and the stimulus-driven salience model (DeepGaze III); rows are the four MultiMatch dimensions. Duration is undefined for the salience model. Young adults do not appear under the adult-prototype reference, for which they are the reference group. Within each panel, the grey “Across all” estimate is the pooled main effect of typicality (from the model with age as a covariate), and the coloured estimates are the per-age-band slopes from the corresponding typicality × age-band interaction model (children, young adults, older adults); because these come from different models, the band slopes need not average onto the pooled estimate. Points are slope estimates with 95% confidence intervals; filled circles denote slopes whose 95% CI excludes zero, open circles those whose CI includes zero. A CI excluding zero indicates a reliable typicality–recognition association within that band but does not by itself imply a difference between bands; age-dependence was tested with the dimension × age-band interaction (Supplementary Table S15).

Directional similarity predicted recognition under every reference (peer OR = 1.11 per s.d., *z* = 3.46, *P* < 0.001; adult OR = 1.15, *z* = 4.16, *P* < 0.001; model OR = 1.21, *z* = 6.31, *P* < 0.001). Duration (peer OR = 1.08, *P* = 0.005 and adult OR = 1.09, *P* = 0.003) and saccade length (peer OR = 1.08, *P* = 0.017) also predicted recognition, whereas positional similarity did not under any reference (all *P* ≥ 0.08) and image memorability had no independent effect (all *P* > 0.50).

We also explored whether the typicality-recognition associations are age-dependent (see details in Supplementary Results). Directional typicality predicted memory broadly across the lifespan, with no age interaction under any reference. Length typicality was adult-emerging — absent in children, reliable in adults (interaction *P* = 0.005 peer, *P* < 0.001 adult). Most strikingly, the relationship of positional typicality to memory reversed with age — positive in children, negative in older adults — but only against human references (interaction *P* = 0.019 peer, *P* = 0.001 adult), not against the model (*P* = 0.072). A per-band partial least-squares analysis was consistent: the memory-related typicality profile was carried by the dynamic dimensions in children and young adults and was not reliable in older adults (Supplementary Results, Supplementary **Fig. S5).**

These effects were not explained by image memorability, which was itself most associated with positional typicality — the one dimension unrelated to recognition — yet the memory-relevant dimensions retained their effects with memorability controlled (Supplementary Table S16). Thus, what predicts memory is conforming in *how* the eyes move, not *where* they are sent; and where spatial conformity does matter, it is human spatial structure that counts.

## Discussion

The central contribution of the present study is a lifespan account of visual exploration that separates where gaze is directed from how it moves through a scene. Decomposing the scanpath into spatial, kinematic and temporal dimensions, and extending the developmental window from early childhood into older adulthood, we find that gaze development is neither uniform across the scanpath properties nor monotonic across age. Spatial targeting was settled early, changed little thereafter, and is well reproduced by a stimulus-driven model. The dynamic structure of viewing — the direction and length of saccades and the duration of fixations — continued to change through childhood, reached maximum convergence in young adulthood, diverged again in later life, and exceeded what a stimulus-driven account could reconstruct. Recognition memory traced the same inverted-U across the lifespan. And when we ask which properties of the encoding scanpath predict what an observer later recognises, the answer converged across every reference frame: typicality in how the eyes moved predicted memory, whereas typicality in where they were sent did not.

At first glance these results seem to pull in different directions and reconciling them is the key to the lifespan pattern. Within-group consistency in the dynamic dimensions follows an inverted-U peaking in young adulthood; the variance decomposition shows the mirror image, with observer-attributable variance highest at both extremes and image-attributable variance peaking in young adulthood. Yet similarity to the young-adult prototype follows no inverted-U — it rises steadily across childhood and is unrelated to age among older adults. These are three views of the same geometry: consistency and observer-attributable variance ask how tightly an age group is packed together, whereas prototype similarity asks how far it sits from a fixed target. That is: children begin both far from the adult pattern and scattered among themselves. Shaped through development, young adults become most alike to each other (more so than any other age groups). Older adults become less alike again, departing in different directions from one another, which is why their mutual consistency falls while their distance from the prototype shows no further gradient across the decade we sampled.

The image-variance trajectory gives this a functional reading. Young adulthood is the age at which gaze is most strongly governed by what each scene affords. At both age extremes, more variance in typicality is carried by stable individual tendencies. Convergence is thus a by-product of stimulus-driven control: when observers track the same images closely they resemble one another, whereas immature and ageing systems track them less closely and personal tendencies fill the gap. This also explains why the effect is confined to the dynamic dimensions. The consistency with which observers direct their eyes to particular locations changes comparatively little across the lifespan — the balance between shared and individual spatial targeting is relatively age-invariant, departing from the common pattern mainly in the youngest children^41,46^, whereas typicality in *how* gaze moves develops for far longer.

This dissociation is one of developmental rate, not absolute alignment: spatial consistency is established early and changes comparatively little thereafter, whereas the consistency of dynamic structure continues to develop well into adulthood. A child may reach adult-like spatial targeting relatively early while still connecting those targets through less organized transitions — remaining within local neighbourhoods, revisiting recently sampled information, or following salient links that do not efficiently traverse the scene’s relational structure. This protracted maturation fits the prolonged development of internally controlled eye movements and the distributed oculomotor network into adolescence^47–49^, whereas basic scene-viewing organisation — ambient-focal alternations, the shortening of fixations and lengthening of saccades — is present from early childhood^22,50^. Notably, these dynamic properties are not independent of one another: because fixation durations follow their own inverted-U across the lifespan, the number of fixations made within a fixed viewing window co-varies with age, so that the developing organisation of where and when the eyes move and how many samples are taken are facets of a single maturing sampling rhythm rather then separable channels. What matures is the efficiency and consistency with which these components are deployed, precisely where the measures operate. In later life the constellation changes again, with increased saccadic latency, altered fixation durations, more local sampling and greater variability^51–53^, alongside accumulated schemas that can guide gaze predictively^54,55^. Older adults may therefore sample under a competition between declining sensory reliability and increasingly influential priors^56^, resolved differently by different individuals — producing the reduced mutual consistency without common age-graded drift that we observe.

Against this background, the memory findings supply the functional link. Across three independent reference frames — age-matched peers, a young-adult prototype and a stimulus-driven model — recognition was better when the encoding scanpath was more typical in its kinematic and temporal organization, and under none when it was more typical in spatial targeting. This argues against an account in which successful encoding depends on fixating a canonical set of scene coordinates^57,58^. The asymmetry follows from the dissociation already described: positional typicality, being dictated by the image, retains little observer-specific variance with which to predict individual differences in memory, whereas the dynamic dimensions carry it. Direction and length index how information is distributed and related across a scene^59^, and fixation duration reflects the balance between processing the current input and initiating the next sample^60,61^; together they determine whether viewing is redundant or efficient and segment it into potentially meaningful encoding events. Position identifies the endpoints of attention; the dynamic dimensions describe the representational structure built between them. The one spatial effect that did emerge was itself age-dependent and human-specific: positional typicality predicted recognition positively in children but negatively in older adults, and only relative to other humans, not the model — so where spatial conformity bears on memory at all, it is conformity to human spatial regularities, not to image salience, that counts.

Two further results substantiate this. Image memorability was most reliably associated with positional canonicality — memorable images drew observers to the same locations — yet position was the one dimension unrelated to recognition. This was not an artifact of statistical control: positional typicality failed to predict recognition whether or not image memorability and image identity were included as covariates. The dimension most tied to the stimulus is thus the one least tied to memory. This converges with the neural literature: eye movements and hippocampal memory processes are closely coordinated during exploration^62–64^, and work combining magnetoencephalography, eye tracking and neural-network modelling found that longer fixations track downstream memory encoding rather than prolonged perceptual processing, with more memorable patches receiving longer fixations and elevated theta–gamma coupling in frontal and hippocampal regions^12^. On that account dwell time indexes encoding engagement rather than difficulty of seeing, consistent with our finding that the typicality of fixation durations predicts recognition — and it also clarifies why duration could be assessed only against human references: the model assigns every fixation the same duration, so it is blind to the dimension most directly tied to encoding. More broadly, this emphasis on sequence over location aligns with the reinstatement literature’s shift from treating the scanpath as part of the stored representation^15^, toward localizing the memory benefit in sequential fidelity over spatial overlap^18,19,65,66^.

The comparison with DeepGaze III sharpens what kind of structure is at stake. The model reproduced positional agreement comparatively well, consistent with much of the spatial consensus in human gaze being recoverable from scene structure and generic sequential dependencies. It reproduced the dynamic dimensions least well — and seeding it with an observer’s own initial fixations did not move its predictions closer to human gaze on any dimension. The missing structure is therefore not a matter of missing initial context the model could be handed, but structure of a kind the model does not represent. The semantic analyses (Supplementary Information) locate part of it concretely: human observers converged most strongly on scenes containing text, a convergence absent in model agents and, developmentally, absent in the youngest children before emerging at reading age — at least partly learned structure, accumulated through experience and expressed most clearly in the temporal control of gaze^1,20^. One caution follows: because DeepGaze III encodes adult, stimulus-driven viewing, lower human–model alignment at the developmental extremes indexes departure from an adult-salience policy, not reduced systematicity, and a model trained on child gaze would plausibly align better with children. Evidence about systematicity comes from within-group consistency, not model alignment; keeping the two distinct is what makes the discrepancy theoretically informative rather than an engineering shortfall, identifying the dimensions in which developmental, physiological and cognitive constraints contribute most to exploration.

A single account accommodates these strands if the scanpath is treated not as a sequence of independently chosen coordinates but as an active sampling policy — decisions about how far to move, in what direction, and when to disengage, taken under uncertainty about what each location will reveal^67^. Such a policy has two separable components. A target function determines where to look; in our data this component was well reproduced by a model trained on image statistics, settled early, and changed little across the lifespan — the profile of a component strongly constrained by the scene. A transport function determines how to move between targets; this component was the one salience model reproduced least well, was not recovered by seeding the model with human fixations, varies between observers at every age, and continued to develop across the lifespan — the profile of a component shaped by more than the image alone. These dynamics are not fixed but adjust to what the observer needs to extract: saccade amplitudes and fixation durations shift with the spatial-frequency and peripheral-visibility properties of a target, whereas saccades that simply continue the previous movement are less sensitive to what it contains — two modes, a default scanning tendency and an information-driven one^61^. Because the memory system receives not a list of coordinates but a temporally structured sequence of samples, it is the transport function that bears on encoding: direction determines which elements are sampled in succession, and therefore which relations are available to be bound, while duration determines how much encoding time each receives, with saccades plausibly acting as boundaries that segment the episode^64^. Consistent with this, gaze transitions following the annotated relational structure of naturalistic scenes occur above chance at encoding and predict subsequent free recall of both object and relational detail, over and above salience and fixation frequency^68^ — and do so at encoding but not retrieval, converging with our finding that it is the organization of sampling during encoding that bears on memory. Typicality is therefore the right measure: the canonical policy is not an arbitrary standard, but the pattern observers converge on when most responsive to the scene, and an observer whose transport function resembles it is sampling in a form the memory system is well suited to encode.

On this view the inverted-U is not a symmetric rise and fall but two processes meeting at a maximum. In childhood the policy is under construction: children possess scene knowledge but implement it through an immature sampling system^47,69–71^, so between-child variation reflects maturation and proximity to the adult pattern indexes developmental progress. In young adulthood the policy has converged, and consistency, stimulus-driven control and recognition peak together. In later life it is displaced rather than regressed: sensorimotor constraint pushes in one direction while accumulated schemas pull in another, and observers balance the two differently. Across both perception and memory, the oldest observers were the least governed by stimulus-intrinsic structure — their gaze diverged most from a salience-driven model, they adjusted least when images constrained gaze more tightly, and their recognition benefited least from intrinsic image memorability. Read this way, gaze is not a record of where attention landed but an observable expression of an active sampling policy — the frame that ties these results together.

### Limitations

Several constraints bound these conclusions. First, the design is cross-sectional, so age differences may reflect cohort effects rather than within-individual change; the gaze-memory associations identify candidate links rather than evidence that altering gaze alters memory, and longitudinal sampling would be needed to trace individual trajectories. Second, our age bands leave adolescence and middle adulthood unmeasured, so the precise form of change between cohorts is unconstrained (orthogonal contrasts and group-midpoint coding gave equivalent results; Supplementary Table S4). Third, our findings rest on a single 60-scene set; on an independent set of 60 content-matched images the trajectory replicated closely (quadratic coefficients correlated r = 0.93 across dimensions, with the young-adult peak recurring for saccade length, direction and fixation duration; Supplementary Note, **Fig. S7)**, the one exception being spatial consistency — which was flat originally but showed weak curvature in replication — indicating that the age-stability of spatial targeting is our least stimulus-general result. Fourth, the gaze–memory effects, although convergent across three reference frames and robust to memorability, are modest and should not be read as a diagnostic marker. Finally, our sample is drawn from a single western, industrialized population, leaving open whether these patterns generalize across cultural contexts — a strong test of the experience-dependent-priors hypothesis. Relatedly, contemporary gaze models are trained on adult-aggregated data and served here as a well-defined null rather than an age-resolved benchmark; fine-tuning them on age-specific data is a natural extension for isolating each age group’s sampling. And because MultiMatch captures the geometry of viewing but not the semantic identity of what was fixated, vision-language-model measures of what was attended — which capture variance partially independent of geometric similarity^72^ — are a complementary dimension future work could add.

### Conclusion

Tracing visual exploration and recognition across the lifespan, we found that the spatial and dynamic properties of the scanpath follow divergent developmental courses with divergent predictive power for memory. Consistency in where the eyes are sent is settled early, conserved across the lifespan, and well captured by a stimulus-driven model, yet carries no advantage for what is later recognised. How the eyes move matures through childhood, peaks in young adulthood, declines in older age, is the aspect of gaze that most exceeds a stimulus-driven account, and is the aspect whose typicality tracks successful encoding. These findings reframe the link between active vision and memory as one of manner rather than location: what is remembered is tied not to the places gaze visits, but to the dynamics with which it moves between them.

## Methods

### Participants

We recruited a lifespan cohort spanning five age groups: children aged 5–7 (n=33), 8-10 (n=39), and 11-12 years (n=31), young adults aged 20–30 years (n=41) and older adults aged 65–80 years (n=35; Supplementary Results for overview; Supplementary Table S1). Participants were recruited from the departmental participant database or by word of mouth. All had normal or corrected-to-normal vision and no history of psychiatric or neurological disorders. Participants, or their legal guardians for minors, provided written informed consent, and were compensated €10 per hour. The study was approved by the ethics committee of the Department of Psychology, Goethe University Frankfurt.

Of 198 participants tested, 195 completed the recognition task. One young adult was excluded as an extreme outlier on recognition sensitivity (d′ above Q3 + 3 × IQR or below Q1 − 3 × IQR^73,74^, leaving 194 for the recognition analysis. Usable eye-tracking data were obtained from 179 participants, who constitute the sample for all scanpaths analyses; of these, 55.20% identified as female and 44.80% as male. Analyses relating gaze to recognition require both measures and were conducted on the 176 participants for whom both were available.

An a priori power analysis (WebPower^74^) indicated that 120 participants would provide power (1−β) =0.95 to detect a between-group effect of f = 0.4 at α = .05, with the effect size based on prior work^75,76^; our sample exceeded this target.

### Apparatus

Stimuli were presented on a 38.1 × 21.4 cm monitor (1,920 × 1,080 pixels, 120Hz) using Psychtoolbox ^77^ and the EyeLink Toolbox in MATLAB R2020b (MathWorks). Images (native 800 × 600 px) were upscaled by a factor of 1.5 to 1,200 × 900 px and presented centrally; at a viewing distance of 46 cm, each image subtended approximately 29.7° × 22.2° of visual angle (40.46 pixels per degree). Eye movements were recorded binocularly with a desktop-mounted EyeLink Portable Duo^78^ at 1,000 Hz running version 6.12 of the Host Software, with head was stabilized on a chin rest. Pupil and corneal-reflection parameters were optimized according to the manufacturer’s recommendations. A 13-point calibration and validation were performed before testing, accepted only when the average error was below 0.5° and the maximum error below 1.0° of visual angle, and drift correction was performed before every trial. Although recording was binocular, the right eye was used for analysis to avoid duplication of fixation and saccade records. Fixations and saccades were defined using the EyeLink default parser^78^.

### Stimuli and Procedure

#### Encoding phase

Participants freely viewed 60 naturalistic scenes for 4 s each while their eye movements were recorded. Stimuli were drawn from the OSIEshort set ^43^, a subset of the OSIE dataset ^44^. Presentation order was held constant across participants to support the investigation of individual differences in viewing.

#### Retrieval phase

Approximately 45 min after encoding, participants completed a cued recognition task. Each trial began with a 1-s probe cue presented under one of six levels of occlusion (30–80%), followed by a 0.5-s mask. Participants then judged whether the cue depicted a studied (“old”) or novel (“new”) scene and rated their recognition confidence on a five-point scale. They were subsequently instructed to reinstate the image by tracing it with their eyes (“Draw the image with your eyes”) for 4 s, after which they provided a second, imagery confidence rating. Probe images comprised the 60 studied scenes and 60 semantic lures; each occlusion level included 10 old and 10 corresponding lure images. Probe presentation order was randomized across participants.

### DeepGaze III scanpath generation

To compare human scanpaths against a stimulus-driven model, we generated synthetic scanpaths with DeepGaze III^41^. Unlike saliency models optimised to predict fixation-density maps, DeepGaze III generates fixation sequences, conditioning each successive fixation on the image and the preceding fixation history.

For each image the model defines a conditional distribution over the next fixation; individual scanpaths are obtained by sampling from it, so repeated runs on the same image yield non-identical sequences. We generated 179 scanpaths per image, each treated as an independent model agent, mirroring the human design of one scanpath per participant per image. Because DeepGaze III is trained on fixations pooled across observers and has no observer-specific parameters, every scanpath is an independent draw from one generative process: the model cannot carry a stable individual signature across images, and inter-agent variability reflects stochastic sampling alone. This variability is the model analogue of observer-level variability in human gaze and provides the baseline for the variance decomposition.

Human scanpaths, acquired at 1,200 × 900 px, were rescaled to the model’s 800 × 600 space for generation, then rescaled back to 1,200 × 900 for similarity computation. The model was initiated with a centre-bias prior matched to the human bias^41^ and a starting fixation drawn from the human ground-truth distribution, and the number of generated fixations was matched to the corresponding human scanpath. In a supplementary seeding analysis, we additionally biased the model with each participant’s first n ∈ {0, 1, 3, 5} fixations before generating the remainder.

### Scanpath similarity

Pairwise scanpath similarity was quantified with the MultiMatch algorithm ^17^, implemented in the authors’ multimatch package. MultiMatch aligns two eye-movement trajectories and compares them along separable, interpretable properties. Of the five dimensions it returns, we analysed four — spatial **Position** (fixation location), saccade **Length** (amplitude), saccade **Direction** (the sequential angular geometry of the scanpath, i.e. its “where-to-next” structure), and fixation **Duration** (dwell timing) — and excluded **Vector**, which combines saccade amplitude and direction into a single composite and is therefore not independent of Length and Direction. This redundancy was confirmed empirically: across all pairwise comparisons Vector correlated *r* = 0.74 with Length, whereas the four analysed dimensions were near-independent of one another (all |*r*| ≤ 0.28; Supplementary Table S2).

All scanpaths were normalised to the stimulus dimensions, and every comparison was computed within image. Similarity was computed for each pair of observers viewing the same scene, excluding self-comparisons, with similarity along each dimension retained separately. Because DeepGaze III assigns every fixation an identical duration (below), the Duration dimension was computed for human–human comparisons only. MultiMatch is a comparative rather than an absolute measure: it does not describe an observer’s viewing in isolation but how closely one trajectory corresponds to another. For every analysis, the focal observer was the unit of aggregation — each observer × scene value is the mean of that observer’s similarity to the relevant set of comparison partners (defined per reference frame, below).

### Reference frames

Scanpath typicality was assessed against three reference frames, each addressing a distinct question. All comparisons were computed within image, and similarity is symmetric, so each pair was included once irrespective of stored ordering, with the focal observer as the unit of aggregation.

#### Peer (within-group) reference

For each participant we computed mean MultiMatch similarity to same-age peers—that is, to all other participants in the same age group on the same image—excluding self-comparisons. This indexes how closely an individual’s scanpath matches the typical viewing pattern of their own age group.

#### Adult-prototype reference

For each participant we computed mean similarity to the young-adult group (20–30 years), excluding self-comparisons; young adults themselves were excluded from this analysis as a focal group (for whom the adult reference would coincide with the peer reference). This indexes alignment with a mature viewing pattern.

#### Model reference

For each participant we computed mean similarity to the full set of DeepGaze III agents. Because model agents carry no age, this comparison was deliberately not age-restricted, in contrast to the age-restricted human references; the number of references scanpaths therefore differs between references, which bears on comparisons of level but not of slope.

### Standardization of similarity scores

Because the four MultiMatch dimensions have different native scales, similarity values were z-standardised before modelling and plotting, using a baseline matched to each question. For *within-group consistency*, each dimention was z-scored against the distribution of all within-group human–human pairs, so zero is the average similarity between two same-aged observers, units are standard deviations (SDs) of that distribution, and positive values indicate above-average within-group agreement. For the *human–model comparison*, the same human–human mean and SD were applied to human–human, human-model, and model-model pairs alike, placing all three on a common, human-anchored scale on which zero is the typical human–human pair and the model similarity is expressed in units of human–human variability. Because the two baselines use different reference distributions, z-scores are comparable within an analysis but not across them.

### Statistical Analysis – general

All analyses were performed in R with lme4, lmerTest, nlme, emmeans and effectsize. LMMs used Satterthwaite degrees of freedom; GLMER coefficients were tested with asymptotic Wald z-tests. Partial effect sizes are reported as ω*^2^_p_*. The significance threshold was α = 0.05 (two-tailed); the multiple-comparison policy is stated per analysis. Age was coded to match each estimand — group rank for the cohort-level variance trajectories, scaled group midpoints for the recognition trajectory and continuous age for the alignment models, and age in months for the child and older-adult models (rationale in Supplementary Methods).

#### Variance Decomposition and Individual Signatures

To quantify how much of scene viewing is governed by the stimulus versus the observer, we decomposed pairwise similarity for each dimension with a linear mixed-effects model containing crossed random intercepts for observer and image and no fixed effects (lme4 ^79^): *similarity ∼ 1 + (1 | observer) + (1 | image)*. Before modelling, pairwise similarities were aggregated to observer level: for each focal observer i and image k, the mean similarity to the other n – 1 partner on that image, excluding self-comparisons (which would inflate similarity and deflate variance). The model partitions total variance into three components: **Agent** variance, the stable idiosyncratic structure an observer carries across scenes (whether an observer is consistently more or less similar to others regardless of what they view); **Image** variance, the extent to which a scene draws all observers into similar viewing; and **Residual** variance, which, given no fixed effects, principally reflects observer-by-image variation, the degree to which an observer’s distinctiveness is scene-specific rather than uniform. Each component is expressed as a proportion of the total (an intraclass correlation, ICC). Whether observer identity carried information was tested by likelihood-ratio comparison against a model omitting the observer term; 95% CIs for variance components were obtained by profile likelihood, robust near the zero boundary.

#### Developmental alignment trajectories

To quantify how children’s scanning matures toward adult and model gaze, we computed for each child one mean similarity score to each reference per MultiMatch dimension, and fitted linear mixed-effects models of similarity on mean-centred age (in months), reference, their interaction and sex, with a by-child random intercept: *similarity ∼ age × reference + sex + (1 | child).* The Age × Reference interaction tested whether the rate of convergence differed between references; per-reference age slopes and reference levels (at mean age) were obtained as estimated marginal trends and means (emmeans) with Tukey-adjusted contrasts. Duration was assessed against the young-adult reference only (DeepGaze III yields no duration metric), using OLS (similarity ∼ age + sex). This analysis included n = 98 children (5–12 years). To test whether convergence on DeepGaze III reflected the model approximating an averaged young adult, the three references (individual young adults, young-adult consensus, DeepGaze III) were entered into a single model per dimension; because the consensus measure showed greater residual variance, reference-specific residual variances were allowed (nlme::lme, varIdent), which improved fit in every dimension (all p < 10⁻¹³). For older adults (65–80; n = 35), the group was referenced to young adults and to DeepGaze III; given the smaller sample, null age effects are read as absence of evidence rather than evidence of stability.

#### Recognition Sensitivity

Recognition sensitivity was quantified with the signal-detection index d′ = Z(hit rate) − Z(false-alarm rate), where a hit was a correct “old” response to a studied scene and a false alarm an “old” response to a lure; extreme rates were handled with the log-linear correction (Hautus, 1995). d′ was computed per participant × occlusion level. We modelled d′ with a linear mixed-effects model including a second-order polynomial for age (to capture the inverted-U trajectory), linear and quadratic terms for occlusion level, image memorability (below) as a z-scored covariate, and a by-participant random intercept. The age-polynomial × occlusion interaction tested whether the developmental trajectory changed under increasing visual challenge. The quadratic occlusion term was retained based on a likelihood-ratio test against the linear-only model, and the threshold structure of the occlusion effect was characterized with Tukey- and Holm-adjusted consecutive-level contrasts (emmeans). A fully categorical age-group × occlusion-level model provided a flexible test of the interaction across all shapes.

#### Image memorability

The intrinsic, image-computable memorability of each scene was estimated with ResMem ^45^, a ResNet-based model predicting human memorability. Memorability scores were included as a covariate (z-scored) in the recognition and gaze–memory models to ensure that effects were not attributable to systematic differences in stimulus memorability. As a design check, the memorability of studied scenes and their lures did not differ (paired t-test, p = 0.51).

#### Gaze typicality predicting recognition

We tested whether the typicality of a participant’s encoding scanpath predicted subsequent recognition of that scene, using generalized linear mixed-effects models (binomial) of trial-level recognition accuracy for studied (old) items. Encoding-phase scanpath similarity to each reference (peer, adult, model), z-scored, served as the predictor, with image memorability and occlusion level as covariates and crossed random intercepts for participant and image: *Correct ∼ Position + Direction + Duration + Length + memorability + occlusion + (1 | participant) + (1 | image).* Models were fitted with the bobyqa optimizer. Age moderation was assessed by adding similarity × age-group interactions, summarized as per-group slopes (emmeans::emtrends). Because the model yields no duration metric, the model-reference analysis omitted Duration and was additionally repeated on low-occlusion trials (≤ 0.5), where the model’s predictions are best validated, to address its training on intact images. Finally, to test whether image memorability could account for gaze–memory associations, we regressed each image’s mean canonical similarity on its memorability; these confound checks are reported uncorrected (the liberal direction for a screen), noting which survive Benjamini–Hochberg FDR correction.

#### Reproducibility

Software: R v4.6.1, lme4 v2.0.1, lmerTest v3.2.1, nlme v3.1.169, emmeans v2.0.3, effectsize v1.0.2, MultiMatch v0.1.3, DeepGaze III v1.1.0, ResMem v1.1.6. Analysis code is available at https://github.com/bhavinc/developmental-scanpaths.

## Acknowledgements

This project was funded by the German Research Foundation (DFG) Research Unit “Abstract Representations in Neural Architectures” FOR 5368 (project number 459426179). It was further supported by the DFG under its Excellence Strategy (EXC 3066/1 “The Adaptive Mind”, Project No. 533717223) and Priority Programme “New Data Spaces for the Social Sciences” (SPP 2431) grant no. 539642788, as well as by the Hessian Ministry of Higher Education, Research, Science and the Arts under the LOEWE Center DYNAMIC (“The Dynamic Network Approach of Mental Health to Stimulate Innovations for Intervention and Change”)

## Notes

https://github.com/bhavinc/developmental-scanpaths

### Competing Interest Statement

The authors have declared no competing interest.

