## Supplementary Information for "How the eyes move, not where they land, predicts what we remember across the lifespan"

Supplementary Methods

The choice of reference

The choice of reference is itself consequential, because any single reference confounds two things at once. Comparing children against adults, as developmental studies typically do, cannot separate immaturity from mere difference from the adult pattern; comparing anyone against a model cannot separate meaningful idiosyncrasy from noise. We therefore assess each scanpath against three references — age-matched peers, a young-adult prototype, and the stimulus-driven model — so that convergent evidence, rather than distance from any one standard, carries the interpretation.

References

***Young-adult consensus control.*** As a developmental control against the possibility that convergence on DeepGaze III merely reflects the model approximating an averaged young adult, we constructed a consensus reference: for each image and dimension we identified the medoid young adult — the individual whose scanpath was, on average, most similar to the remaining young adults — and computed each participant's similarity to this representative scanpath.

**Coding of age across analyses**.

Age enters three analyses on three codings, each matched to its estimand, and we state the difference explicitly to avoid confusion. (i) The pooled and age-resolved variance trajectories are fitted to five cohort-level estimates and code age as age-group rank (1–5); midpoint coding would place the older-adult cohort far from the others (scaled midpoint z = 2.96 against −0.56 to 0.45 for the remaining groups) and let a single cohort dominate the quadratic term. (ii) The observation-level trajectory analyses (recognition; alignment) use scaled group midpoints or continuous age, as noted. (iii) The child and older-adult alignment models use chronological age in months. Because the cohorts are unequally spaced, trajectory shape was additionally characterised with orthogonal polynomial contrasts on the ordered age-group factor, which decompose group means into linear and quadratic components without assuming spacing between cohorts or interpolating across unsampled ages; group-midpoint coding gave equivalent results.

**Comparability of denominators.**

Because similarity is pairwise, averaging over n partners shrinks the residual by ≈ 1/n while leaving Agent and Image variance unaffected; since ICCs are ratios, every component's share depends on n. Humans and model agents were therefore averaged over comparable numbers of partners so that their decompositions sit on the same scale. To compare the magnitude of individuality directly, we report the ratio of human to DeepGaze III agent variance (infinite where the model produced a singular fit with zero agent variance) and the percentage-point difference in image-driven variance (Δ Image = Var_human − Var_DG).

**Age as a nuisance in the pooled fit.**

Because the sample spans 5–79 years, pooled Agent variance could in principle reflect systematic age-group differences rather than individual idiosyncrasy. Each model was therefore refitted with age group as a fixed effect; the resulting Agent estimates index individual structure over and above developmental stage.

**Age-resolved variance decomposition**

To examine how the composition of gaze variance changes across the lifespan, the decomposition above was repeated separately within each age group and dimension. For this analysis, similarity was computed among same-age observers only — within-group similarity is the appropriate reference for a developmental comparison — and these estimates are compared only across age groups, never against the model.

Within each of the 20 cells (5 age groups × 4 dimensions), Agent and Image components were estimated by bootstrapping over observers (1,000 iterations): observers were resampled with replacement (rows within an observer are not independent) and the decomposition refitted to each resample, yielding a mean ICC, its bootstrap standard error, and a 95% percentile CI per cell. Whether observer identity carried information within a cell was tested by likelihood-ratio comparison against a model omitting the observer term.

Developmental trajectories were then fitted to these cohort-level estimates. For each component we fitted a weighted linear mixed model with a quadratic effect of age and a random intercept for dimension.

ICC ~ poly(age, 2) + (1 | dimension), weights = 1 / SE²

Each estimate was weighted by the inverse of its squared bootstrap standard error, so more precisely estimated cells contributed more; this weighting reflects estimation uncertainty in each ICC and is distinct from the between-observer variance the ICC itself quantifies. The random intercept accounts for the non-independence of the four dimensions, which share observers and images. Age was coded as age-group rank (1–5) for the reason given above.

Because the three components are proportions of one total, they sum to one in every cell and their trajectories are not independent: a decline in one is necessarily offset by a rise in the others. The Agent and Image trajectories therefore describe a single reallocation rather than two independent effects and are reported as such (the linear coefficients sum to ≈ 0 as required; the quadratic coefficients sum to ≈ 0 up to differences in weighting and shrinkage between the separately fitted models).

Two further tests were computed per dimension: whether each component's trajectory departed from linearity (quadratic term of a weighted polynomial regression), and whether Agent and Image followed different trajectories (Component × age interaction, tested by model comparison; residual df = 4). These answer different questions and can diverge — a component may curve reliably while not curving differently from its counterpart. Both are exploratory: with five cohorts these dimension-wise tests have limited power, and the composite models are the primary analysis.

**Partial least squares correlation (PLSC).**

As a multivariate complement to the trial-level models, we used behavioural partial least squares correlation to test whether the joint profile of scanpath typicality across MultiMatch dimensions covaried with recognition, and whether that covariation differed across the lifespan. PLSC was run separately within each age band (children, young adults, older adults) and for each reference template (peer, young-adult prototype, salience model), on subject-level data (one value per participant per dimension, averaged over trials).

For each analysis, the typicality block X (the z-scored MultiMatch dimensions — Position, Direction, Length and, for the human references, Duration) and the memory variable Y (mean recognition accuracy) were first residualised on age in months within the band, so that the resulting latent variable reflected viewing strategy rather than maturation, and then standardised. Because Y was univariate, the analysis reduced to the singular value decomposition of the between-block correlation vector R = Xᵀy; the singular value quantifies the overall covariation between the typicality profile and memory, and the corresponding left saliences give each dimension's contribution to that covariation.

Statistical significance of the singular value was assessed by permutation (5,000 permutations), randomly reordering Y relative to X and recomputing the singular value to build a null distribution; P is the proportion of permuted singular values equalling or exceeding the observed value. The stability of each dimension's salience was assessed by bootstrap resampling of participants (5,000 resamples); a bootstrap ratio (salience divided by its bootstrap standard error) with absolute value greater than 2 (approximately P < 0.05) was taken to indicate a reliable contribution. Analyses were conducted in R [version]; the adult-prototype reference excludes the young-adult band (its reference group) and the salience-model reference excludes Duration (undefined for the model).

**Scene category classification**

Scene categories were assigned automatically using a multi-detector pipeline implemented in MATLAB (R2023b, Computer Vision Toolbox and Deep Learning Toolbox), combining object detection, face detection, and language-agnostic text detection. Each of the 60 scenes was processed independently.

*Object detection*. Objects were detected with a YOLOv4 network (CSP-DarkNet-53 backbone) pretrained on the COCO dataset. Detections were retained at a confidence threshold of 0.25; if no detection survived this threshold, a relaxed threshold of 0.10 was applied to avoid classifying sparsely populated scenes as empty. Because people are the category most consequential for our contrasts, an additional lenient pass retained "person" detections at a confidence of 0.05 when the standard thresholds returned none. Detected COCO labels were grouped a priori into animals (bird, cat, dog, horse, sheep, cow, elephant, bear, zebra, giraffe), food and tableware (banana, apple, sandwich, orange, broccoli, carrot, hot dog, pizza, donut, cake, bowl, cup, bottle), vehicles (bicycle, car, motorcycle, airplane, bus, train, truck, boat), technology (laptop, cell phone, TV, keyboard, mouse, remote), and household or manipulable objects (book, clock, vase, scissors, teddy bear, toothbrush, chair, couch, bed, dining table).

*Face detection*. Faces were detected with Viola–Jones cascade classifiers (frontal and profile models) in a two-pass procedure. The first pass used a strict merge threshold (8) and discarded detections smaller than max(30 px, 4% of the shorter image dimension). If no face survived, a second pass relaxed the merge threshold (4), upscaled the image by a factor of 1.6 to recover small faces, and applied a correspondingly smaller minimum size (max(24 px, 3% of the shorter dimension)). This two-pass design was adopted to reduce misses for distant or partially occluded faces while limiting false positives in the primary pass.

*Text detection.* Text was detected with CRAFT (Character Region Awareness for Text detection), a language-agnostic detector that localizes text regions without recognizing characters, ensuring that detection did not depend on the language of the depicted text. Images larger than 1,024 px on their longest side were downscaled prior to detection for numerical stability, and detected regions were rescaled to original coordinates. Regions smaller than 0.015% of image area, or narrower than 12 px in either dimension, were discarded as spurious. An image was classified as containing text if at least two regions survived filtering; requiring two rather than one region reduced false positives from isolated high-contrast edges.

*Classification fallback.* When object detection returned no labels at any threshold, a pretrained image-classification network (ResNet-18) provided a rescue pass: the top 20 class labels with confidence ≥ 0.02 were examined, and keyword matches to food or technology terms set the corresponding category flag. This fallback was applied only when object detection was empty and did not override positive detections.

*Category assignment.* Each image received three binary indicators: object presence (any animal, food, technology, or household-object detection), human presence (any person detection or any detected face), and text presence. Images were then assigned to a single category for analysis, with text taking precedence: any image containing detected text was classified as TEXT; among remaining images, those containing objects were classified as OBJECT; those containing people but no objects as HUMAN; and images with no detection in any channel as OTHER (excluded from category analyses). A composite tag recording all detected content types (e.g. OBJ+HUM) was retained for each image to document category overlap.

Supplementary Results

Participants

Intellectual ability was assessed with the RIAS. Because the Verbal (VIX), Nonverbal (NIX) and General (GIX) indices are age-normed (population M = 100, SD = 15), each score indexes a participant's standing relative to their own age norm. All groups scored near the normative mean (Supplementary Table Sx). Age groups did not differ in general intellectual ability (GIX: F(4, 144) = 0.18, P = 0.95, η² = 0.005) or verbal ability (VIX, Welch's F(4, 66.7) = 1.53, P = 0.20, η² = 0.04). A weak difference in nonverbal ability (NIX: F(4, 145) = 2.74, P = 0.031, η² = 0.07) did not survive correction for the three indices tested (Holm-adjusted P = 0.093) or a variance-robust test (Welch's P = 0.053), and the effect was small. Overall, the age samples were comparable in normed intellectual ability, indicating that the reported age differences in gaze and memory are unlikely to reflect ability differences between cohorts.

Supplementary Table S1. Participant characteristics by age group.

| **Characteristic** | **5–7** | **8–10** | **11–12** | **18–30** | **65–80** | **Overall** |
| --- | --- | --- | --- | --- | --- | --- |
| **N** | 33 | 40 | 31 | 41 | 35 | 180 |
| **Female, n (%)** | 19 (58%) | 17 (42%) | 18 (58%) | 28 (68%) | 16 (46%) | 98 (54%) |
| **Age, M (SD)** | 6.3 (0.6) | 9.0 (0.8) | 11.5 (0.6) | 22.8 (2.6) | 71.3 (3.6) | 24.2 (24.0) |
| **Age, range** | 5–7 | 8–10 | 11–13 | 19–29 | 65–79 | 5–79 |
| **Right-handed, n (%)** | 27 (82%) | 32 (80%) | 27 (87%) | 31 (76%) | 32 (94%) | 149 (83%) |
| **Corrected vision, n (%)** | 2 (6%) | 6 (15%) | 3 (10%) | 9 (22%) | 28 (80%) | 48 (27%) |
| **RIAS GIX, M (SD)** | 111.9 (11.2) | 109.5 (23.2) | 110.7 (10.7) | 108.8 (9.0) | 109.7 (9.1) | 109.9 (14.1) |
| **RIAS VIX, M (SD)** | 113.7 (13.2) | 108.1 (14.3) | 110.2 (10.7) | 108.8 (7.7) | 105.8 (10.1) | 108.9 (11.5) |
| **RIAS NIX, M (SD)** | 107.2 (10.5) | 103.2 (11.9) | 108.3 (9.7) | 106.5 (10.0) | 111.6 (10.2) | 107.2 (10.8) |

*Note:* Values are n (%) for categorical variables and mean (SD) for continuous variables; age range in years. RIAS Verbal (VIX), Nonverbal (NIX) and General (GIX) indices are age-normed (population mean 100, SD 15); [n] denotes the number with a valid score, as RIAS was not obtained for all participants. Corrected vision = wore correction during testing.

Supplementary Table 2. Pearson correlations between MultiMatch dimensions, computed across all pairwise observer comparisons.

| **Dimension** | **Vector** | **Length** | **Position** | **Direction** | **Duration** |
| --- | --- | --- | --- | --- | --- |
| Vector | 1.00 |  |  |  |  |
| Length | 0.74 | 1.00 |  |  |  |
| Position | 0.33 | 0.28 | 1.00 |  |  |
| Direction | 0.25 | 0.09 | 0.22 | 1.00 |  |
| Duration | 0.07 | 0.09 | 0.07 | 0.08 | 1.00 |

**Natural scene gaze separates into a stimulus-driven and an individual component**

These signatures were not reducible to developmental stage, including age group as a fixed effect reduced agent variance by no more than 1.7 percentage points on any dimension, indicating that the idiosyncratic structure an observer carries across scenes is largely independent of their age.

**Gaze composition follows developmental trajectory**

Stable individual differences were present in every age group and on every dimension (likelihood ratio tests against models omitting the observer term, all P < 0.001

Dimension-specific analyses showed this reallocation to be uneven (**Fig. 2d**): Agent and Image followed reliably different age trajectories for Direction (component × age interaction, F_(2,4)_ = 13.3, P = 0.017) and Duration (F_(2,4)_ = 10.1, P = 0.027), and marginally so for Length and Position (P = 0.062 and 0.064). Component-wise, idiosyncrasy curved reliably only for Direction (quadratic P = 0.014) and stimulus-dependency only for Duration (P = 0.045).

**Supplementary Table 3.** Variance decomposition of scanpath similarity.

| **Dimension** | **Source** | **Agent (%)** | **Image (%)** | **Residual (%)** | **LRT chi2** | **P** |
| --- | --- | --- | --- | --- | --- | --- |
| Length | Human | 18.35 [14.84, 22.98] | 28.06 [19.93, 41.19] | 53.59 | 2561.4 | < 0.001 |
|  | DG III | 0.07 [0.00, 0.22] | 66.73 [47.56, 97.72] | 33.20 | 1.3 | 0.263 |
| Position | Human | 12.63 [10.17, 15.89] | 36.26 [25.79, 53.19] | 51.10 | 1782.9 | < 0.001 |
|  | DG III | 0.06 [0.00, 0.15] | 79.82 [56.91, 116.83] | 20.12 | 2.4 | 0.123 |
| Direction | Human | 14.46 [11.63, 18.19] | 22.10 [15.66, 32.51] | 63.44 | 1685.6 | < 0.001 |
|  | DG III | 0.00 [0.00, 0.15] | 49.75 [35.41, 72.92] | 50.25 | 0.0 | 1.000 |
| Duration | Human | 16.40 [13.18, 20.65] | 7.57 [5.28, 11.28] | 76.03 | 1595.0 | < 0.001 |

*Note:* Agent (%) is the proportion of total variance attributable to observer identity (the individual signature); Image (%) the proportion attributable to the stimulus; Residual (%) the remainder, which principally reflects observer-by-image variation — the extent to which an observer's distinctiveness is specific to particular scenes rather than uniform across them. Brackets give 95% bootstrap percentile confidence intervals (1,000 iterations, resampling observers). LRT compares models with and without the observer term; near-zero values for DeepGaze III reflect the absence of observer-specific parameters in the model. Fixation duration is undefined for DeepGaze III, which assigns every fixation an identical duration. DG III, DeepGaze III; LRT, likelihood-ratio test; χ², chi-squared.

Supplementary Table S4. Human variance components with and without adjustment for age group.

| **Dimension** | **Agent,**  **unadjusted (%)** | **Agent,**  **age-adjusted (%)** | **Difference (pp)** | **Image (%)** | **Residual (%)** |
| --- | --- | --- | --- | --- | --- |
| Length | 18.30 | 17.48 | 0.82 | 28.45 | 54.07 |
| Position | 12.58 | 12.01 | 0.58 | 36.55 | 51.45 |
| Direction | 14.58 | 13.27 | 1.30 | 22.21 | 64.52 |
| Duration | 17.31 | 15.58 | 1.73 | 7.80 | 76.62 |

*Note.* Unadjusted values are from the models reported in Table 1; adjusted values are from the same models refitted with age group as a fixed effect, so that Agent variance indexes individual structure over and above developmental stage. Image and Residual columns are from the age-adjusted models. N = 181 observers, 60 scenes.

Supplementary Table S5. Likelihood-ratio tests for the observer (Agent) term within each age group and MultiMatch dimension, comparing the crossed-random-effects model with and without a random intercept for observer.

| **Dimension** | **Age group** | **chi2** | **df** | **P** |
| --- | --- | --- | --- | --- |
| Length | 5-7 | 537.0 | 1 | 8.4e-119 |
|  | 8-10 | 168.8 | 1 | 1.4e-38 |
|  | 11-12 | 413.6 | 1 | 5.9e-92 |
|  | 20-30 | 243.7 | 1 | 6.1e-55 |
|  | 65-80 | 754.8 | 1 | 3.6e-166 |
| Position | 5-7 | 650.4 | 1 | 1.8e-143 |
|  | 8-10 | 110.9 | 1 | 6.1e-26 |
|  | 11-12 | 118.7 | 1 | 1.2e-27 |
|  | 20-30 | 184.9 | 1 | 4.1e-42 |
|  | 65-80 | 381.4 | 1 | 6.1e-85 |
| Direction | 5-7 | 318.4 | 1 | 3.2e-71 |
|  | 8-10 | 65.3 | 1 | 6.6e-16 |
|  | 11-12 | 21.8 | 1 | 3.0e-06 |
|  | 20-30 | 56.2 | 1 | 6.5e-14 |
|  | 65-80 | 457.9 | 1 | 1.4e-101 |
| Duration | 5-7 | 187.7 | 1 | 1.0e-42 |
|  | 8-10 | 220.0 | 1 | 9.2e-50 |
|  | 11-12 | 110.2 | 1 | 8.7e-26 |
|  | 20-30 | 142.7 | 1 | 6.9e-33 |
|  | 65-80 | 433.9 | 1 | 2.2e-96 |

Note: A significant test indicates that observer identity carries reliable information within that cell.

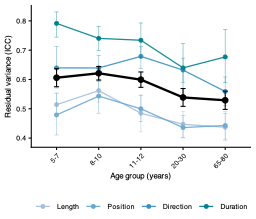

**Supplementary Figure S1** | Residual variance across the lifespan. Residual variance — the component of scanpath similarity attributable to neither observer identity nor image, principally reflecting observer-by-image variation — as a function of age group. Coloured lines show each dimension; the black line shows the mean across dimensions. Error bars are 95% bootstrap confidence intervals (1,000 resamples of observers) for dimensions, and pooled bootstrap standard errors for the composite. Residual variance declined linearly with age in the composite model (b = −0.16, t = −4.86, P < 0.001; quadratic P = 0.131); per-dimension trends were negative in all four dimensions but individually unreliable (all P > 0.09). Agent and Image components are shown in Fig. 2c,d. N = 181 observers, 60 scenes.

Supplementary Table S6. Shape of the age-related change in within-group scanpath consistency, per MultiMatch dimension.

| **Dimension** | **Test** | **Estimate** | **SE** | **Stat** | **P** |
| --- | --- | --- | --- | --- | --- |
| Length | Fall (20-30 - 65-80) | 0.003 | 0.002 | 1.93 | 0.055 |
|  | Peak vs extremes | 0.007 | 0.001 | 4.51 | 2.4e-05 |
|  | Rise (20-30 - 5-7) | 0.010 | 0.002 | 5.72 | 1.4e-07 |
|  | Trend: linear | 0.018 | 0.004 | 4.58 | 9.0e-06 |
|  | Trend: quadratic | -0.014 | 0.005 | -2.88 | 0.004 |
| Position | Fall (20-30 - 65-80) | -0.009 | 0.004 | -2.45 | 0.046 |
|  | Peak vs extremes | -0.002 | 0.003 | -0.56 | 0.575 |
|  | Rise (20-30 - 5-7) | 0.005 | 0.004 | 1.46 | 0.291 |
|  | Trend: linear | 0.027 | 0.008 | 3.16 | 0.002 |
|  | Trend: quadratic | -0.000 | 0.010 | -0.00 | 0.999 |
| Direction | Fall (20-30 - 65-80) | 0.014 | 0.003 | 4.47 | 1.8e-05 |
|  | Peak vs extremes | 0.014 | 0.003 | 5.30 | 1.0e-06 |
|  | Rise (20-30 - 5-7) | 0.014 | 0.003 | 4.57 | 1.8e-05 |
|  | Trend: linear | 0.004 | 0.007 | 0.58 | 0.564 |
|  | Trend: quadratic | -0.056 | 0.009 | -6.48 | 9.0e-10 |
| Duration | Fall (20-30 - 65-80) | 0.036 | 0.005 | 6.70 | 5.5e-10 |
|  | Peak vs extremes | 0.034 | 0.005 | 7.37 | 2.0e-11 |
|  | Rise (20-30 - 5-7) | 0.032 | 0.005 | 5.87 | 2.2e-08 |
|  | Trend: linear | 0.016 | 0.012 | 1.31 | 0.192 |
|  | Trend: quadratic | -0.115 | 0.015 | -7.75 | 7.4e-13 |

Note: Trend rows give the linear and quadratic components of the five ordered age-group means (orthogonal polynomial contrasts). Contrast rows test whether the young-adult group exceeds the two age extremes (Peak vs extremes) and whether consistency rises from the youngest children to young adults and falls again in older adults (Rise, Fall). All from linear mixed-effects models with crossed random intercepts for observer and image; P values Holm-adjusted within dimension for the contrasts.

Supplementary Table S7. Pairwise contrasts of within-group scanpath consistency between age groups, per MultiMatch dimension (Tukey-adjusted; linear mixed-effects models with crossed random intercepts for observer and image).

| **Dimension** | **Comparison** | **Estimate** | **SE** | **t** | **P** |
| --- | --- | --- | --- | --- | --- |
| Length | 11-12 vs 20-30 | -0.004 | 0.002 | -2.20 | 0.185 |
|  | 11-12 vs 65-80 | -0.001 | 0.002 | -0.32 | 0.998 |
|  | 20-30 vs 65-80 | 0.003 | 0.002 | 1.93 | 0.304 |
|  | 5-7 vs 11-12 | -0.006 | 0.002 | -3.25 | 0.012 |
|  | 5-7 vs 20-30 | -0.010 | 0.002 | -5.72 | < 0.001 |
|  | 5-7 vs 65-80 | -0.007 | 0.002 | -3.68 | 0.003 |
|  | 5-7 vs 8-10 | -0.005 | 0.002 | -2.86 | 0.038 |
|  | 8-10 vs 11-12 | -0.001 | 0.002 | -0.57 | 0.979 |
|  | 8-10 vs 20-30 | -0.005 | 0.002 | -2.96 | 0.029 |
|  | 8-10 vs 65-80 | -0.002 | 0.002 | -0.93 | 0.886 |
| Position | 11-12 vs 20-30 | 0.003 | 0.004 | 0.67 | 0.962 |
|  | 11-12 vs 65-80 | -0.006 | 0.004 | -1.64 | 0.476 |
|  | 20-30 vs 65-80 | -0.009 | 0.004 | -2.45 | 0.107 |
|  | 5-7 vs 11-12 | -0.008 | 0.004 | -2.01 | 0.266 |
|  | 5-7 vs 20-30 | -0.005 | 0.004 | -1.46 | 0.589 |
|  | 5-7 vs 65-80 | -0.014 | 0.004 | -3.73 | 0.002 |
|  | 5-7 vs 8-10 | -0.007 | 0.004 | -1.97 | 0.284 |
|  | 8-10 vs 11-12 | -0.001 | 0.004 | -0.15 | 1.000 |
|  | 8-10 vs 20-30 | 0.002 | 0.004 | 0.56 | 0.981 |
|  | 8-10 vs 65-80 | -0.007 | 0.004 | -1.89 | 0.327 |
| Direction | 11-12 vs 20-30 | 0.001 | 0.003 | 0.44 | 0.992 |
|  | 11-12 vs 65-80 | 0.015 | 0.003 | 4.59 | < 0.001 |
|  | 20-30 vs 65-80 | 0.014 | 0.003 | 4.47 | < 0.001 |
|  | 5-7 vs 11-12 | -0.016 | 0.003 | -4.69 | < 0.001 |
|  | 5-7 vs 20-30 | -0.014 | 0.003 | -4.57 | < 0.001 |
|  | 5-7 vs 65-80 | -0.001 | 0.003 | -0.17 | 1.000 |
|  | 5-7 vs 8-10 | -0.011 | 0.003 | -3.55 | 0.004 |
|  | 8-10 vs 11-12 | -0.004 | 0.003 | -1.38 | 0.642 |
|  | 8-10 vs 20-30 | -0.003 | 0.003 | -1.02 | 0.847 |
|  | 8-10 vs 65-80 | 0.011 | 0.003 | 3.44 | 0.007 |
| Duration | 11-12 vs 20-30 | 0.002 | 0.005 | 0.42 | 0.993 |
|  | 11-12 vs 65-80 | 0.038 | 0.006 | 6.66 | < 0.001 |
|  | 20-30 vs 65-80 | 0.036 | 0.005 | 6.70 | < 0.001 |
|  | 5-7 vs 11-12 | -0.034 | 0.006 | -5.89 | < 0.001 |
|  | 5-7 vs 20-30 | -0.032 | 0.005 | -5.87 | < 0.001 |
|  | 5-7 vs 65-80 | 0.004 | 0.006 | 0.70 | 0.956 |
|  | 5-7 vs 8-10 | -0.008 | 0.005 | -1.40 | 0.626 |
|  | 8-10 vs 11-12 | -0.026 | 0.006 | -4.74 | < 0.001 |
|  | 8-10 vs 20-30 | -0.024 | 0.005 | -4.65 | < 0.001 |
|  | 8-10 vs 65-80 | 0.012 | 0.005 | 2.16 | 0.201 |

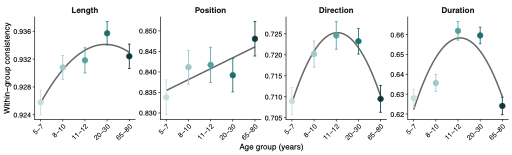

**Supplementary Figure S2 | Developmental trajectory of within-group scanpath consistency, by dimension.** Estimated marginal means of within-group similarity for each age group (points; ±1 s.e.m.), from linear mixed-effects models with crossed random intercepts for observer and image. Curves show the fitted trend across the ordered age groups: a quadratic function for Length, Direction and Duration, and a linear function for Position, following the significant components identified by orthogonal polynomial contrasts (Supplementary Table Sx). The x-axis is ordinal (age groups equally spaced by rank), so the curves describe the pattern across sampled cohorts and do not interpolate across unsampled ages. Direction and Duration follow a symmetric inverted-U peaking in young adulthood (quadratic P = 9.0 × 10⁻¹⁰ and 7.4 × 10⁻¹³); Length rises and largely plateaus (linear P = 9.0 × 10⁻⁶, quadratic P = 0.004); Position increases monotonically (linear P = 0.002, quadratic P = 0.999).

**Fixation count does not account for the developmental consistency trajectories**

Because average fixation duration follows an inverted-U across the lifespan, the number of fixations made within the fixed 4-s encoding window differs modestly with age (F(4, 174) = 5.31, P < 0.001, η² = 0.11; means ranging from 11.7 fixations in the youngest children to 13.4 in young adults), and fixation count is itself a strong correlate of within-group similarity on the dynamic dimensions (Duration, Direction, Length: all P < 10⁻⁸⁰) though not of positional similarity (P = 0.96). We therefore refitted each dimension's age-group model with each observer's fixation count included as a covariate. The developmental effects were retained on every dimension, with only modest attenuation of the count-coupled dimensions (Duration ω²_p 0.31 → 0.25; Direction 0.18 → 0.17; Length 0.14 → 0.12; Position unchanged at 0.06; all age effects P < 10⁻⁴). The age differences in within-group consistency are thus not reducible to age differences in the number of fixations made.

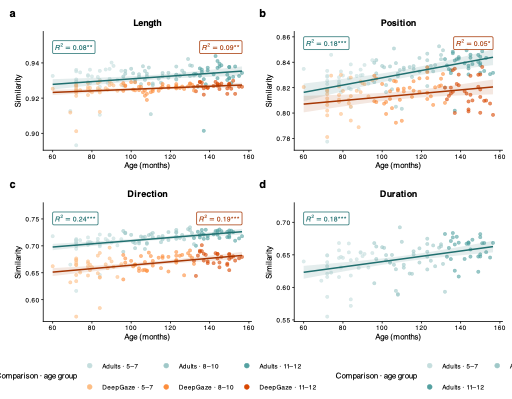

**Figure S3 | Developmental convergence of children's gaze toward human-adult and model references.** Overlaid scatter plots show each child's mean scanpath similarity to two references — the young-adult consensus (reference cohort 20–30 years) and the DeepGaze III ensemble — as a function of age in months. Panels correspond to MultiMatch dimensions arranged by functional category: saccadic amplitude (Length), spatial target selection (Position), saccadic grammar (Direction), and fixation timing (Duration); Duration is shown for the young-adult reference only, as DeepGaze III provides no duration metric. Each point is one child's mean similarity across all stimuli; point colour encodes reference × age cohort — similarity to young adults in teal (5–7 light, 8–10 medium, 11–12 dark) and similarity to DeepGaze III in orange (same age ordering; see legend). Coloured lines are linear regression fits (teal = young-adult reference, orange = DeepGaze III) with shaded 95% confidence intervals; colour-matched labels report adjusted R² and significance for each reference. Similarity increased with age for both references across all metrics, but the rate of convergence on the young-adult consensus significantly exceeded that on DeepGaze III in every dimension (Age × Reference interaction; see Results), most markedly for Position, where the model slope did not differ from zero. n = 98 children (both references). Significance: *p < 0.05, **p < 0.01, ***p < 0.001.

Supplementary Table S8. Test of whether DeepGaze III is reducible to an averaged adult. For each MultiMatch dimension, children's convergence slope (change in similarity per month) toward three references: individual young adults, the per-image consensus (medoid) young adult, and DeepGaze III. The consensus adult is the young adult whose scanpath is, per image and dimension, most similar to all other young adults (the most representative adult observer). Models used reference-specific residual variances (nlme::lme with varIdent); the two rightmost columns give the Tukey-adjusted slope contrasts. Fixation duration is excluded (undefined for the model).

| **Dimension** | **Young adults** | **YA consensus** | **DeepGaze III** | **Young adults - DeepGaze III** | **YA consensus - DeepGaze III** |
| --- | --- | --- | --- | --- | --- |
| Length | 7.57e-05  (t=2.45, 0.017) | 5.15e-05  (t=1.51, 0.136) | 4.27e-05  (t=1.09, 0.278) | 3.30e-05  (t=1.38, 0.354) | 8.76e-06  (t=0.31, 0.947) |
| Position | 2.98e-04  (t=4.02, 1.2e-04) | 4.14e-04  (t=4.97, 3.5e-06) | 1.84e-04  (t=2.31, 0.023) | 1.13e-04  (t=3.81, 5.7e-04) | 2.30e-04  (t=4.74, 1.3e-05) |
| Direction | 3.04e-04 (  t=4.86, 5.4e-06) | 4.32e-04  (t=5.52, 3.7e-07) | 3.31e-04  (t=4.41, 3.0e-05) | -2.71e-05  (t=-0.66, 0.789) | 1.02e-04  (t=1.62, 0.241) |

**The model is not reducible to an averaged adult.** Comparing convergence slopes across three references — individual young adults, a per-image consensus (medoid) adult, and DeepGaze III — children approached the consensus adult faster than the model for Position (slope difference t(188) = 5.77, P < 0.001), so on this dimension the model does not capture the adult spatial pattern even at the population level. For saccade length the consensus and model slopes were indistinguishable (t(193) = 0.38, P = 0.92): for this low-level property DeepGaze III is equivalent to an averaged young adult. For direction the difference was marginal (t(191) = 2.14, P = 0.084). Individual young adults exceeded the consensus for Position and Direction but not Length, consistent with children's gaze approaching specific rather than prototypical observers.

Supplementary Table S9. Developmental convergence of children's scanpaths on young adults and on DeepGaze III. Slopes are the change in similarity per month (age centred), from linear mixed models (Similarity ~ age x reference + sex + (1|child)), Satterthwaite df. The difference column is the reference x age interaction. Fixation duration is undefined for DeepGaze III.

| **Dimension** | **Adult slope** | **Model slope** | **Difference (adult - model)** |
| --- | --- | --- | --- |
| Length | 7.57e-05 (t=3.07, 0.003) | 4.27e-05 (t=1.73, 0.086) | 3.30e-05 (t=1.38, 0.172) |
| Position | 2.98e-04 (t=4.25, 5.2e-05) | 1.84e-04 (t=2.63, 0.010) | 1.13e-04 (t=3.81, 2.6e-04) |
| Direction | 3.04e-04 (t=4.32, 3.6e-05) | 3.31e-04 (t=4.71, 8.0e-06) | -2.71e-05 (t=-0.66, 0.514) |
| Duration | 4.13e-04 (t=4.85, 4.6e-06) | — (model has no duration) | — |

Supplementary Table S10. Older adults (65-80 y): similarity to the young-adult prototype and to DeepGaze III showed no relation to age, and the two references did not differ in age slope, on any dimension (all P values shown). Same model as the child analysis (Similarity ~ age x reference + sex + (1|participant)). N = 32 older adults.

| **Dimension** | **Adult slope (p)** | **Model slope (p)** | **Interaction (p)** |
| --- | --- | --- | --- |
| Length | 0.781 | 0.858 | 0.883 |
| Position | 0.821 | 0.708 | 0.734 |
| Direction | 0.743 | 0.505 | 0.483 |

**Saccade amplitude does not account for the seeding age effects.**

To test whether the age differences in seeding reflected oculomotor rather than strategic differences, we computed saccade amplitudes from the encoding fixations. Amplitude followed an inverted-U across the lifespan (quadratic β = −0.53, P = 0.021, R² = 0.24; group-level quadratic P < 10⁻⁸), peaking in childhood and young adulthood and declining in older age (older adults below every younger group, Tukey all P < 0.001; no differences among younger groups). Horizontal saccade bias showed no lifespan trend. Two features rule out a motor account of the seeding effects. Children's amplitudes matched young adults', so the child seeding effects cannot stem from immature saccades; and older adults, whose amplitudes were furthest from the adult-trained model's scale, showed the smallest length penalty — the opposite of what motor misalignment would predict. The age effect in seeding is therefore spatial, not oculomotor.

**Seeding**

Critically, this amplitude analysis rules out a simple motor account of the seeding effects rather than supplying one. Because children's saccade amplitudes matched those of young adults, the child-related seeding effects cannot stem from immature saccade amplitude—addressing a potential developmental confound. And although older adults made shorter saccades, this cannot explain their smaller Length penalty: their amplitude was the furthest of any group from the adult-trained model's scale, the opposite of what a motor-alignment account would predict. The interpretable signal is spatial. Older adults' fixated locations — not the metric properties of their saccades —are what the model most fails to reproduce, and this divergence concerns where the eyes are directed rather than how far they move.

Supplementary Table S11. Quadratic age term for recognition sensitivity (d′), fitted with continuous age (primary) and with group midpoints (robustness). Both show a significant negative quadratic (inverted-U). Occlusion and image memorability controlled; random intercept for participant.

| **Coding** | **Quadratic b** | **t** | **P** |
| --- | --- | --- | --- |
| Continuous age (primary) | -4.05 | -3.15 | 0.002 |
| Continuous age (primary) | -8.98 | -7.03 | 3.4e-11 |
| Continuous age (primary) | 7.33 | 2.17 | 0.030 |
| Continuous age (primary) | 1.83 | 0.54 | 0.587 |
| Group midpoints (robustness) | -3.69 | -2.85 | 0.005 |
| Group midpoints (robustness) | -8.68 | -6.74 | 1.6e-10 |
| Group midpoints (robustness) | 6.81 | 2.01 | 0.044 |
| Group midpoints (robustness) | 0.05 | 0.02 | 0.988 |

Supplementary Table S12. Consecutive-step contrasts of d′ between adjacent occlusion levels (Holm-adjusted), marginal over age. Recognition was preserved across mild degradation and fell in discrete steps at higher occlusion.

| **Step** | **Estimate** | **SE** | **Stat** | **P** |
| --- | --- | --- | --- | --- |
| OcclusionLevel0.4 - OcclusionLevel0.3 | -0.242 | 0.234 | -1.03 | 0.906 |
| OcclusionLevel0.5 - OcclusionLevel0.4 | 0.056 | 0.119 | 0.47 | 0.906 |
| OcclusionLevel0.6 - OcclusionLevel0.5 | -0.448 | 0.058 | -7.72 | 1.4e-13 |
| OcclusionLevel0.7 - OcclusionLevel0.6 | -0.079 | 0.090 | -0.88 | 0.906 |
| OcclusionLevel0.8 - OcclusionLevel0.7 | -0.409 | 0.076 | -5.41 | 3.3e-07 |

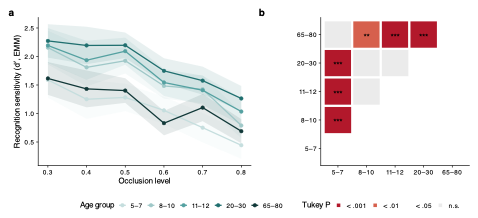

**Figure 4 |**  **a,** Model-estimated d′ (EMM) across six occlusion levels, one line per age group. Recognition was preserved across mild-to-moderate occlusion (0.3–0.5) and declined in discrete steps thereafter; the age groups declined in parallel, with no age × occlusion interaction. **b**, Tukey-adjusted pairwise comparisons of d′ between age groups (averaged over occlusion); cell color indicates the significance level and asterisks the p-value bin.

Supplementary Table S13. Pairwise differences in d′ between age groups (Tukey-adjusted), occlusion and memorability controlled. Positive = first-named group higher. Localises the inverted-U.

| **Comparison** | **Estimate** | **SE** | **Stat** | **P** |
| --- | --- | --- | --- | --- |
| 5-7 - 8-10 | -0.537 | 0.114 | -4.71 | 4.4e-05 |
| 5-7 - 11-12 | -0.639 | 0.113 | -5.66 | 4.3e-07 |
| 5-7 - 20-30 | -0.814 | 0.113 | -7.19 | 1.3e-10 |
| 5-7 - 65-80 | -0.117 | 0.111 | -1.05 | 0.832 |
| 8-10 - 11-12 | -0.103 | 0.113 | -0.91 | 0.893 |
| 8-10 - 20-30 | -0.277 | 0.110 | -2.53 | 0.089 |
| 8-10 - 65-80 | 0.420 | 0.108 | 3.89 | 0.001 |
| 11-12 - 20-30 | -0.174 | 0.112 | -1.55 | 0.531 |
| 11-12 - 65-80 | 0.523 | 0.110 | 4.73 | 4.2e-05 |
| 20-30 - 65-80 | 0.697 | 0.107 | 6.51 | 6.9e-09 |

Supplementary Table S14. Memorability x age interaction in trial-level recognition of old items (GLMM, logit). Coefficients are the memorability slope for each age group relative to the youngest (5-7). Older adults show a significantly weaker memorability benefit.

| **Term** | **b (log-odds)** | **z** | **P** |
| --- | --- | --- | --- |
| memorability (5-7 ref) | 0.10 | 0.91 | 0.362 |
| memorability x 8_10 | -0.04 | -0.58 | 0.563 |
| memorability x 11_12 | 0.04 | 0.57 | 0.572 |
| memorability x 20_30 | 0.03 | 0.34 | 0.732 |
| memorability x 65_80 | -0.20 | -2.79 | 0.005 |

**Memorability**

Target and lure images did not differ in predicted memorability (paired t(59) = −0.66, p = .51), confirming that recognition differences between conditions are not attributable to a memorability imbalance. Memorability was related to gaze typicality, but the pattern did not track the memory effects. Its strongest and most consistent association was with positional canonicality — under all three references (peer β= 0.16, model β = 0.22, adult β = 0.18; all FDR q = 0.044) — the one dimension that predicted recognition under no template. Memorability was also associated with directional and, for the human references, duration typicality, but more weakly (direction: peer β = 0.13, adult β = 0.13, both q = 0.044; duration: peer β = −0.08, q = 0.044), and no association reached significance for length (all q > 0.9). Critically, every gaze–memory effect was estimated with image memorability and image identity controlled, so the directional, length, and duration effects on recognition hold over and above any covariation between memorability and gaze (Fig. 7b). Conforming to a reference's exploration thus predicts encoding independent of how memorable the studied image is.

Supplementary Table S15. Dimension x age-band interaction tests (Type III Wald chi-square) for the typicality x age models. A significant interaction licenses interpretation of the per-band slopes in Supplementary Table S3; reliable interactions were obtained for Length and Position under the human references. The adult-prototype models contrast children and older adults only (young adults are the reference group), hence df = 1.

| **Template** | **Dimension** | **chi^2** | **df** | **P** |
| --- | --- | --- | --- | --- |
| Adult | Direction | 0.80 | 1 | 0.371 |
|  | Duration | 0.11 | 1 | 0.746 |
|  | Length | 16.06 | 1 | 6.2e-05 |
|  | Position | 10.65 | 1 | 0.001 |
| Model | Direction | 3.26 | 2 | 0.196 |
|  | Length | 5.53 | 2 | 0.063 |
|  | Position | 5.27 | 2 | 0.072 |
| Peer | Direction | 0.86 | 2 | 0.651 |
|  | Duration | 4.01 | 2 | 0.135 |
|  | Length | 10.67 | 2 | 0.005 |
|  | Position | 7.94 | 2 | 0.019 |

**Supplementary Results — Multivariate typicality profile and memory (PLSC)**

To test whether the multivariate profile of scanpath typicality — rather than each dimension considered separately — relates to recognition, we ran behavioural partial least squares correlation (PLSC) within each age band and reference template, with age (in months) residualised from both blocks so that the latent variable reflects viewing strategy rather than maturation (Methods). This treats the correlated MultiMatch dimensions jointly rather than as competing predictors and provides a multivariate check on the trial-level models.

The typicality profile covaried reliably with memory in children under the adult-prototype reference (singular value 0.40, P_perm = 0.023) and suggestively under the peer reference (P_perm = 0.079), and in young adults the profile showed the richest structure (peer reference, latent-variable–memory r = 0.29). Across these bands the covariation was carried by the dynamic dimensions: Direction, saccade Length and — in young adults — fixation Duration all contributed reliably (bootstrap ratio |BSR| > 2), whereas positional typicality carried no reliable salience in any band or template (all |BSR| < 1.6). Length was the most consistent contributor, reaching reliability in children under every template (peer, adult and salience-model references). In older adults, no typicality profile related reliably to memory under any reference (all P_perm > 0.31); the profile saliences in this group were negative in sign — the reverse valence from children and young adults — but sat within non-significant latent variables and are therefore interpreted only as consistent with the sign reversal seen in the trial-level Position analysis, not as an independent effect.

The PLSC thus converges with the trial-level models: what covaries with memory is the dynamic organisation of viewing (direction, length, timing), not where the eyes are sent, with the effect strongest in children and young adults and unreliable in the older-adult sample.

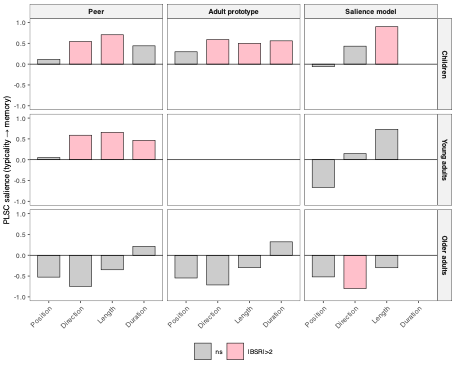

**Supplementary Fig. S5 , Multivariate typicality–memory profile by age band and reference template (PLSC).** Bootstrap-derived saliences (contribution of each MultiMatch dimension to the typicality–memory latent variable) from behavioural PLSC run separately within children, young adults and older adults (rows) for the peer, adult-prototype and salience-model references (columns). Age in months was residualised from both blocks within each band before analysis. Bars are salience weights; solid fill denotes reliable contributions (|bootstrap ratio| > 2 across 5,000 resamples), grey denotes non-reliable. Duration is undefined for the salience-model reference. Young adults do not appear under the adult-prototype reference, for which they are the reference group. Positional typicality contributes reliably in no band; the memory-related profile is carried by the dynamic dimensions. Older-adult latent variables were not reliable (all permutation P > 0.31; 5,000 permutations) and are shown for completeness. Saliences are the weights of each dimension in the typicality–memory latent variable (unit-length vector; sign = direction of the memory relationship, magnitude = relative contribution); they are not independent effect sizes.

Supplementary Table S16. Gaze typicality and recognition: odds ratios (per +1 s.d. similarity) for each MultiMatch dimension under three reference templates, from mixed-effects logistic models controlling for image memorability, occlusion and age group, with random intercepts for participant and image. Duration is undefined for the salience model.

| **Template** | **Dimension** | **OR** | **95% CI** | **z** | **P** |
| --- | --- | --- | --- | --- | --- |
| Peer | Direction | 1.10 | [1.04, 1.17] | 3.46 | 5.4e-04 |
|  | Duration | 1.08 | [1.02, 1.14] | 2.84 | 0.005 |
|  | Length | 1.08 | [1.01, 1.15] | 2.40 | 0.017 |
|  | Position | 1.05 | [0.98, 1.12] | 1.35 | 0.176 |
| Adult prototype | Direction | 1.15 | [1.08, 1.23] | 4.16 | 3.1e-05 |
|  | Duration | 1.09 | [1.03, 1.16] | 2.97 | 0.003 |
|  | Length | 1.00 | [0.93, 1.08] | 0.04 | 0.966 |
|  | Position | 1.07 | [0.99, 1.16] | 1.74 | 0.082 |
| Salience model | Direction | 1.21 | [1.14, 1.28] | 6.31 | 2.8e-10 |
|  | Length | 1.05 | [0.98, 1.13] | 1.41 | 0.158 |
|  | Position | 1.01 | [0.95, 1.07] | 0.40 | 0.687 |

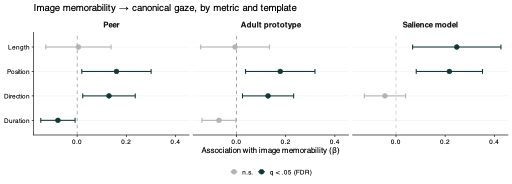

**Fig. 6 |**. **b,** Memorability–gaze associations: standardised regression coefficients (points) with 95% CIs (bars) for the relationship between image memorability (ResMem) and the canonical (template-typical) value of each scanpath metric. Dashed line, β = 0; points coloured by FDR-corrected significance (q < .05); error bars are nominal 95% CIs. Memorability was related to gaze typicality most consistently for spatial position—the one dimension that did not predict recognition (a)—whereas the memory-relevant dimensions were weakly or inconsistently associated with memorability.

Supplementary Table S17. Image memorability and false alarms to lures, by age group. Per-group slopes (log-odds of a false alarm per +1 s.d. of the lure image's memorability) from a binomial mixed model of false-alarm responses to lure (new) items, with centred occlusion and crossed random intercepts for participant and image. No group's false alarms were reliably predicted by memorability; in particular, older adults' false alarms were unrelated to memorability (slope 0.01, P = 0.92), indicating that their reduced old-item memorability benefit does not reflect heightened false recognition of memorable-looking lures.

| **Age group** | **Slope (log-odds/s.d.)** | **95% CI** | **z** | **P** |
| --- | --- | --- | --- | --- |
| 5–7 | -0.051 | [-0.266, 0.164] | -0.47 | 0.642 |
| 8–10 | 0.037 | [-0.178, 0.253] | 0.34 | 0.735 |
| 11–12 | 0.221 | [-0.012, 0.454] | 1.86 | 0.064 |
| 20–30 | 0.018 | [-0.210, 0.246] | 0.15 | 0.878 |
| 65–80 | 0.010 | [-0.193, 0.214] | 0.10 | 0.920 |

**Gaze typicality and recognition**

**Main effects.**

Encoding-scanpath typicality predicted subsequent recognition of studied items, with the pattern consistent across all three references (Supplementary Table S1). Directional typicality was the most robust predictor (peer OR = 1.11 per s.d., z = 3.46, P < 0.001; adult OR = 1.15, z = 4.16, P < 10⁻⁴; model OR = 1.21, z = 6.31, P < 10⁻⁹). Duration predicted recognition under the peer (OR = 1.08, z = 2.84, P = 0.005) and adult (OR = 1.09, z = 2.97, P = 0.003) references, and saccade length under the peer reference (OR = 1.08, z = 2.40, P = 0.017). Positional typicality predicted recognition under no reference (peer P = 0.18, adult P = 0.08, model P = 0.69), and image memorability had no independent effect in any model (all P > 0.5).

**Age-dependence.**

We decomposed these effects across three age bands (children, young adults, older adults), testing dimension × age-band interactions (Supplementary Table S2) before interpreting per-band slopes (Supplementary Table S3). Directional typicality showed no reliable interaction with age under any reference (peer χ²(2) = 0.86, P = 0.65; adult χ²(1) = 0.80, P = 0.37; model χ²(2) = 3.26, P = 0.20); its per-band slopes were positive throughout and reliable in children and older adults under all references, indicating an age-general benefit rather than a group-specific one. Length typicality showed a reliable interaction under both human references (peer χ²(2) = 10.67, P = 0.005; adult χ²(1) = 16.06, P < 10⁻⁴): its slope was null in children (peer b = 0.00 [−0.08, 0.08]) but positive in young adults (b = 0.22 [0.09, 0.36]) and older adults (b = 0.16 [0.04, 0.29]), an adult-emerging effect. Duration typicality benefited young adults specifically (peer b = 0.19 [0.06, 0.31]), with children and older adults null.

Positional typicality showed the clearest age-dependent pattern: a reliable interaction under both human references (peer χ²(2) = 7.94, P = 0.019; adult χ²(1) = 10.65, P = 0.001) but not against the model (χ²(2) = 5.27, P = 0.072). Its slope was positive in children (peer b = 0.10 [0.03, 0.18]; adult b = 0.12 [0.04, 0.20]) and negative in older adults (peer b = −0.10 [−0.24, 0.04]; adult b = −0.15 [−0.30, 0.00]), a reversal in valence across the lifespan. Against the salience model no band showed a reliable positional effect (all P > 0.09), and the interaction was not significant — so the positional reversal is specific to typicality measured relative to other humans, not to stimulus salience.

**Memorability does not account for the associations.**

Every gaze–memory effect above was estimated with image memorability controlled. To test the confound directly, we regressed each image's mean canonical typicality on its memorability, per dimension and reference (Supplementary Table S4). Memorability was most consistently associated with positional canonicality — the one dimension unrelated to recognition — and least with the dynamic dimensions that predicted memory; associations reported uncorrected, with Benjamini–Hochberg FDR-corrected values in Supplementary Table S4. The memory-relevant dimensions retained their effects with memorability partialled out.

**Replication across an independent stimulus set**

Participants returned after approximately two weeks and free-viewed an independent set of 60 images under identical conditions. Because task, instructions and observers were held constant while the stimuli changed entirely, agreement between sessions indexes the stability of age-related gaze structure across both stimuli and time, rather than measurement reliability alone.

To test whether the developmental patterns reflect properties of viewing behaviour rather than the specific images used, we repeated the core analyses on an independent set of 60 images. These served as recognition lures and were matched to the original set in semantic category and content but shared no visual overlap with it, so replication cannot be attributed to the particular stimuli that generated the main results. Of the 179 participants in the main analysis, 156 also viewed the independent set; this is therefore a within-sample test of stimulus generalization rather than replication in a new sample.

The group-level similarity structure replicated. Across all 25 cells of the 5 × 5 age-group similarity matrix, the two stimulus sets correlated strongly for Direction (r = 0.91) and Length (r = 0.76), and moderately for Position (r = 0.63) and Duration (r = 0.57; all P < 0.01). The relative similarity of each age group to every other was thus largely a property of how groups view scenes rather than of the images viewed.

The developmental trajectory replicated in magnitude. The quadratic age coefficients of the two stimulus sets correlated strongly across dimensions (r = 0.93, P < 0.001): the dimensions with the steepest inverted-U in the main data — Duration and Direction — again showed the largest negative quadratic terms in the independent set (Direction β = −0.035; Duration β = −0.049; both P < 0.001), and Length again showed a shallower inverted-U (β = −0.009, P < 0.001). The relative ordering of developmental curvature across dimensions was therefore preserved on entirely different images.

One dimension diverged. Positional similarity, which showed no reliable curvature in the main data (β = 0.001, P = 0.90), showed a modest inverted-U in the independent set (β = −0.019, P < 0.001). The monotonic profile of spatial target selection was thus not fully robust across stimulus sets: on the content-matched images, spatial consistency also peaked weakly in young adulthood rather than rising monotonically. Because Position carried the weakest developmental effect in both sets, this divergence concerns the shape of a small effect rather than its magnitude, but it indicates that the manner-versus-location dissociation is clearest for the strongly developing dimensions (Direction, Duration) and less sharp for spatial targeting.

The semantic-category organization replicated. The within-group advantage — greater gaze similarity among same-age peers — and its ordering across text, object and human scenes correlated between stimulus sets (r = 0.86, P < 0.001), indicating that the category structure of gaze similarity generalizes beyond the specific images.

Overall, the developmental trajectory, its ordering across dimensions, and the semantic-category structure of gaze reproduced on an independent, content-matched image set, confirming that the main patterns are properties of age-related viewing behaviour rather than of the specific stimuli. The one exception — a weak inverted-U for spatial position where the main data showed none — suggests the monotonic character of spatial targeting is the least stimulus-general aspect of the results.

The developmental peak reproduced across stimulus sets: for Length, Direction and Duration, within-group similarity peaked at young adults (20–30 years) in both the original and the independent image set. Only Position diverged — its trajectory was flat in the original set (no meaningful peak) and weakly peaked at young adults in the independent set — consistent with spatial targeting being the one dimension whose developmental shape did not generalize across stimuli.

Replication held at the level of individual age groups. Discrepancies in age-group similarity were small and scattered (largest cell difference 0.18 z units, with no age group systematically divergent), and the developmental peak fell at young adults (20–30) in both stimulus sets for Length, Direction and Duration. Position again formed the exception: flat in the original set and weakly peaked in the independent set, it was the only dimension whose peak location did not match — reinforcing that spatial targeting is the least developmentally and stimulus-generalizable dimension of gaze.

At the level of individual age groups, the two stimulus sets agreed closely: the largest discrepancy in any age-group × dimension cell was 0.18 z units, and discrepancies were small and scattered across groups and dimensions rather than concentrated in any one group, indicating no systematic failure to replicate at particular ages. (Per-group correlations across the four dimensions are uninformative given only four values per group and are not reported.)

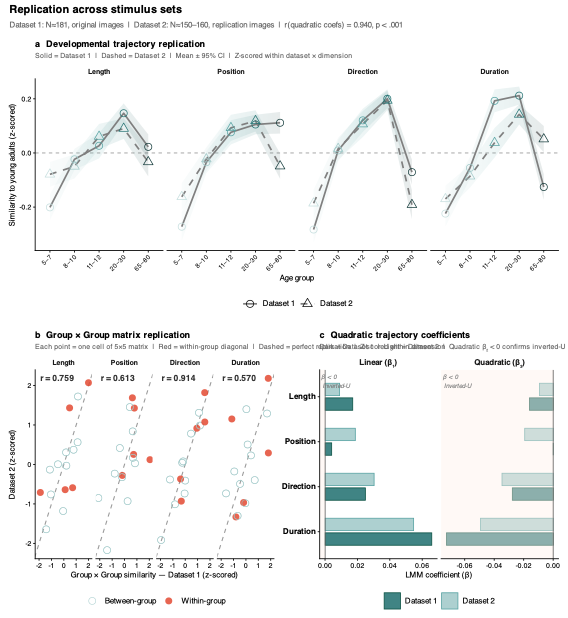

**Supplementary Figure 7 | Replication of developmental gaze patterns across an independent, content-matched stimulus set.** Comparison of the main analysis (Dataset 1: 60 original images, *N* = 179) with an independent replication set (Dataset 2: 60 content-matched lure images sharing no visual overlap with the originals; *N* = 156, all also in Dataset 1). **a**, Developmental trajectory of within-group similarity (z-scored, mean ± 95% CI) across age groups, per MultiMatch dimension, for both stimulus sets. **b**, Agreement of the full 5 × 5 age-group similarity matrix between sets; each point is one matrix cell (red, within-group diagonal), z-scored within dimension; the dashed line marks perfect replication, and *r* is the per-dimension cross-set correlation. **c**, Linear and quadratic age coefficients from the trajectory models for both sets; negative quadratic terms (shaded region) indicate an inverted-U. The quadratic coefficients correlated *r* = 0.93 (*P* < 0.001) across dimensions. Direction and Duration reproduced a clear inverted-U on both sets; Position, monotonic in Dataset 1, showed a weak inverted-U in Dataset 2 (see Supplementary Note X).

**Semantic category structure in human but not model gaze**

Scenes differ in the semantic priors they invoke: text imposes a learned scanning routine, faces and people attract socially-guided attention, and object scenes offer neither. We asked whether gaze similarity varies by scene category, and whether any such structure exceeds what a stimulus-driven model reproduces. Images were classified as containing text, people, or objects, and we compared within-group human similarity with model–model similarity, which indexes how strongly the image itself constrains scanpaths.

Human gaze showed reliable category structure, concentrated in the temporal dimension: within-group similarity differed by category most strongly for fixation duration (F(2, 57) = 9.41, P < 0.001, ω²p = 0.22) and modestly for saccade length (F(2, 57) = 3.50, P = 0.037, ω²p = 0.08), with no reliable effect for position or direction. The model showed no category structure on any dimension (all P ≥ 0.06; direction F(2, 57) = 0.02, P = 0.98, ω²p = 0.00), indicating that the image properties DeepGaze III encodes do not themselves differentiate these scene categories.

The category profile therefore differed between human observers and model agents (category × source interaction: Length F(2, 20305) = 653, ω²p = 0.06; Position F = 112, ω²p = 0.01; Direction F = 18.7, ω²p = 0.002; all P < 10⁻⁸). The divergence was driven by text scenes: humans were substantially more similar to one another than model agents were for text images (Length −0.43 z, Position −0.15 z), whereas for object and people scenes the model agents were as similar or more so (Length +0.47 and +0.24 z respectively). Shared human scanning of text thus exceeds the convergence attributable to image structure alone.

This text-specific convergence showed a developmental signature consistent with reading acquisition. For saccade length, the youngest children showed no text advantage over object scenes (5–7: Δ = 0.0006, P = 0.97), whereas every older group did (8–10: Δ = −0.007, P = 0.025; 11–12: Δ = −0.007, P = 0.026; 20–30: Δ = −0.009, P = 0.004). For fixation duration the text advantage was present from the youngest group but strengthened into young adulthood (5–7: Δ = −0.016; 20–30: Δ = −0.024) before weakening in older adults (Δ = −0.010, P = 0.13). The kinematic signature of text-driven scanning therefore appears to emerge with reading fluency rather than being present from early childhood.

These category × age interactions were statistically reliable but small in absolute terms (ω²p = 0.002–0.015), reflecting the large number of observations rather than large effects; the category main effects, estimated at the image level, are the more interpretable quantities. With approximately 20 images per category, these analyses should be treated as exploratory. Because the replication set was always viewed in the second session, stimulus and session order are confounded; the modest differences we observe in positional similarity could in principle reflect either the images or increased familiarity with the procedure.
